# Alpha-synuclein regulates nucleolar DNA double-strand break repair in melanoma

**DOI:** 10.1101/2024.01.13.575526

**Authors:** Moriah R. Arnold, Gabriel M. Cohn, Kezia Catharina Oxe, Somarr N. Elliott, Cynthia Moore, Peter V. Laraia, Sahar Shekoohi, Dillon Brownell, Charles K. Meshul, Stephan N. Witt, Dorthe H. Larsen, Vivek K. Unni

## Abstract

Although an increased risk of the skin cancer melanoma in people with Parkinson’s Disease (PD) has been shown in multiple studies, the mechanisms involved are poorly understood, but increased expression of the PD-associated protein alpha-synuclein (αSyn) in melanoma cells may be important. Our previous work suggests that αSyn can facilitate DNA double-strand break (DSB) repair, promoting genomic stability. We now show that αSyn is preferentially enriched within the nucleolus in the SK-MEL28 melanoma cell line, where it colocalizes with DNA damage markers and DSBs. Inducing DSBs specifically within nucleolar ribosomal DNA (rDNA) increases αSyn levels near sites of damage. αSyn knockout increases DNA damage within the nucleolus at baseline, after specific rDNA DSB induction, and prolongs the rate of recovery from this induced damage. αSyn is important downstream of ATM signaling to facilitate 53BP1 recruitment to DSBs, reducing micronuclei formation and promoting cellular proliferation, migration, and invasion.

## Introduction

Neurodegeneration and cancer are traditionally thought of as opposite processes, with the former characterized by neuronal death and the latter uncontrolled cellular proliferation. In general, neurodegenerative disease patients are at moderately decreased risk of developing most cancers^1^, however exceptions exist. The best described positive association between a neurodegenerative disease and cancer risk is between Parkinson’s Disease (PD) and the skin cancer melanoma^2^. Since the initial observation by Skibba et al.^3^, many epidemiological studies report an increase in melanoma risk ranging from 1.4- to 20-fold among individuals with PD compared to healthy individuals^1, 4-19^. More malignant forms of melanoma that originate in the head or neck region^20^ are also associated with sporadic and genetic forms of PD^21^. This risk is bidirectional, since there is also increased risk of developing PD in melanoma patients (1.7-4.2-fold)^11, 14, 22-24^. Altogether, the association between PD and melanoma is well-established, yet the cause is poorly understood.

One potential molecular link is the neurodegeneration-associated protein alpha-synuclein (αSyn) that aggregates into cytoplasmic inclusions called Lewy bodies in PD and other Lewy body disorders (LBDs)^25,26^. This Lewy pathology is associated with neuronal death in midbrain dopaminergic neurons, cortical glutamatergic neurons, and other vulnerable cell populations in LBDs^27^. αSyn is not only abundant in the central nervous system, but is also found in other cell types in the body, including skin^28, 29^. Given this connection between PD and melanoma, the presence of αSyn in the skin is intriguing. αSyn is not highly expressed in melanocytes of healthy individuals, but increased in melanocytes of people with PD^30, 31^, and in ∼85% of primary and metastatic melanoma samples^4, 31-33^. Many immortalized human melanoma cell lines also show high expression of αSyn^1, 4, 32, 34^. These findings suggest that αSyn upregulation may be a key molecular link between PD and melanoma and an important contributor in both disease pathologies.

Our previous work using longitudinal *in vivo* multiphoton imaging in mice revealed that Lewy body formation in cortical neurons was associated with a loss of soluble αSyn from the nucleus and cytoplasm and that only neurons bearing Lewy bodies went on to die^35^. This led us to hypothesize that loss of a normal nuclear αSyn function could contribute to these cells’ demise. We unexpectedly found that αSyn colocalized with DNA double-strand break (DSB) repair components within the nucleus of human and mouse cells. Alpha-synuclein knockout (KO) increased DSBs and impaired DSB repair efficiency in human cells and mouse cortical neurons, which could be rescued by transgenic reintroduction of human αSyn in the mouse neuron αSyn KO background^36^. Our previous findings are consistent with the work of others observing activation of DNA damage response (DDR) pathways in synucleinopathy models^37-39^. We suggested that in neurodegenerative disease αSyn loss-of-function within the nucleus of inclusion-bearing neurons may contribute to their cell death^36^. We also speculated that an αSyn gain-of-function in melanoma, where nuclear αSyn is upregulated, could play a potentially protective role against DNA damage. Consistent with this, work by others has shown that αSyn KO human melanoma cells implanted as xenografts in mice exhibit slower growth and increased apoptosis^40^, paired with reduced tumor-induced mechanical allodynia^41^. In addition, WT melanoma cells in αSyn overexpressing mice show increased metastasis^42^, indicating a potentially complex interaction between αSyn within melanoma cells and other tissues in the body. Alpha-synuclein expression is also correlated with poorer survival for patients with melanoma^40^. Although numerous studies have described localization of αSyn to the nucleus in melanoma cells^31, 34, 43-46^, its nuclear function is unclear. In order to better understand these interesting links between neurodegeneration and cancer, we set out to test the function of αSyn within the nucleus of melanoma cells.

## Results

### Alpha-synuclein localizes to the nucleolus

Unexpectedly, our immunocytochemical staining showed clear enrichment of αSyn within the nucleolar sub-compartment of the nucleus in melanocyte and melanoma cells. In the SK-Mel28 melanoma cell line, αSyn colocalized to DAPI-poor regions and with established nucleolar markers treacle, nucleophosmin, and nucleostemin (Figure 1A, 1B). Similar results were also seen in A375 (melanoma line), PIG1 (melanocyte line), and human primary melanocyte cells derived from foreskin (Supplemental Figure 1). We next used fluorescence deconvolution analysis to measure αSyn localization at higher spatial resolution within nucleolar sub-compartments. Mammalian nucleoli are comprised of three sub-compartments: the fibrillar center, the dense fibrillar component, and the granular component (rRNA transcription occurs at the interface of the FC and DFC, early rRNA processing occurs in the DFC, followed by additional rRNA processing and pre-ribosome assembly in the GC). These sub-compartments were visualized using antibodies specific for each of them (RPA194, fibrillar center; fibrillarin, dense fibrillar component; nucleophosmin, granular component). 3D modeling (Imaris) revealed the localization of αSyn primarily in the granular component, directly adjacent to the dense fibrillar component, and relatively excluded from fibrillar centers (Figure 1C). In order to obtain more detailed spatial information, we measured αSyn localization with immunogold transmission electron microscopy (immuno-TEM). Immuno-TEM images in SK-Mel28 cells also showed particle labeling primarily in the granular component and/or dense fibrillar component, but relatively excluded from the fibrillar center compared to an αSyn KO cell line previously described^40^ (Figure 1D, Supplemental Figure 2). Taken together, these data strongly suggest that αSyn localizes to the nucleolus in melanocytes and melanoma cells, and is enriched within the granular component of the nucleolus. These data are consistent with other work showing αSyn labelling within the nucleolus^34, 44^, and extends it by demonstrating preferential enrichment within the granular component of the nucleolus.

**Figure 1.**
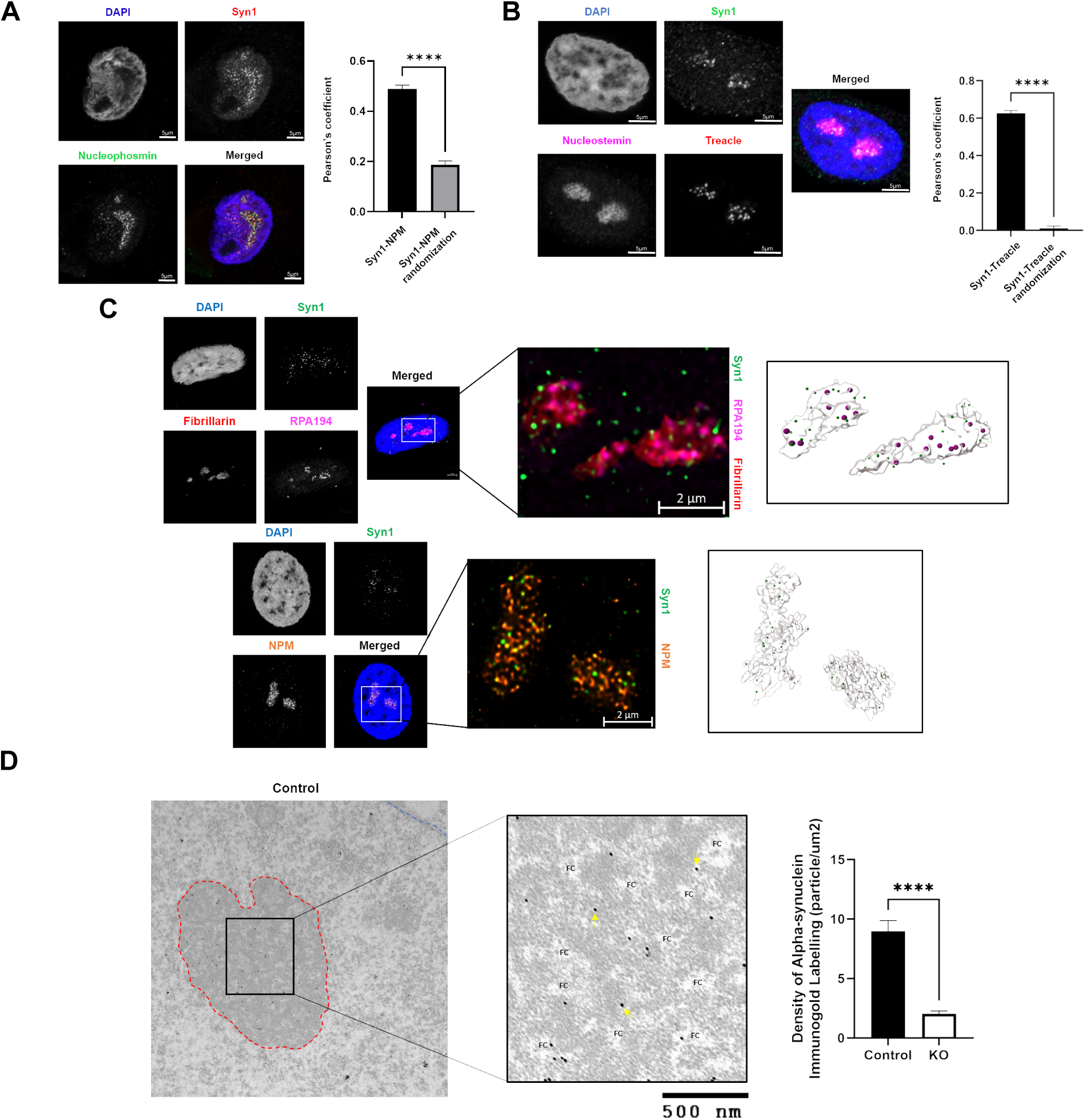
Alpha-synuclein localizes to the granular component of the nucleolus. A, B) SK-Mel28 cells were seeded on PLL-coated coverslips and then fixed and stained for alpha-synuclein (Syn1), nucleolar markers (nucleophosmin, treacle, and nucleostemin), and DAPI. Cells were imaged on the Zeiss 980 confocal microscope with Airyscan and colocalization was analyzed in Imaris software. Error bars represent Standard Error of the Mean (SEM) with quantification from 3 biological replicates. ****p<0.0001 by T-test. C). SK-Mel28 cells were seeded on PLL-coated coverslips and then fixed and stained for alpha-synuclein (Syn1), fibrillar center marker (RPA194), dense fibrillar component marker (fibrillarin), granular component marker (nucleophosmin), and DAPI. Cells were imaged on the Zeiss 980 confocal microscope with Airyscan oversampling and Joint Deconvolution and Channel Alignment post-processing. 3D renderings were produced using Imaris software. D) SK-Mel28 control and KO cell pellets were fixed with 0.1M sodium cacodylate buffer (pH 7.2) containing 0.05% glutaraldehyde, 4% paraformaldehyde, and 0.1% picric acid for 2 hours at RT. Thin sections were cut on an ultramicrotome (EM UC7) using a diamond knife. Sections were permeabilized and stained for alpha-synuclein (MJFR1, 1:75) (12nm colloidal gold particles). The stained sections were observed by a JEOL 1400 transmission electron microscope. Red labelling denotes nuclear membrane. Blue labelling denotes the outline of the nucleolus. FC=Fibrillar Center. Arrows point to representative MJFR1 staining. Quantification of 3 biological replicates. ****p<0.0001 by T-test. Error bars denote SEM.

### Alpha-synuclein knock-out increases γH2AX levels in the nucleus and nucleolus in an ATM- and ATR-dependent manner

In order to understand the functional role of αSyn enrichment within the nucleolus, we measured levels of the phosphorylated histone marker of DSB repair γH2AX within the nucleoplasmic and nucleolar compartments independently. αSyn and γH2AX levels were both significantly greater in the nucleolus compared to the nucleoplasm and this γH2AX strongly colocalized with αSyn foci in SK-Mel28 cells (Figure 2A). We also measured significant colocalization between αSyn and γH2AX in PIG1 (melanocyte cell line) and primary melanocytes (Supplemental Figure 3A, 3B). PLA analysis revealed significant αSyn and γH2AX colocalization, within the ∼40nm resolution (Figure 2B), suggesting close proximity of these two molecules in nuclear and nucleolar DSB repair-associated foci. Interestingly, αSyn KO cells exhibited increased γH2AX immunocytochemical signal in the nucleus and this difference was even larger in the nucleolus (Figure 2C), suggesting that αSyn is even more important in the nucleolus for DSB repair than it is in the nucleoplasm. Importantly, stable reintroduction of wild-type human αSyn into the αSyn KO background (αSyn KI-rescue) reduced both nucleolar and nuclear γH2AX back to control levels (Figure 2C, Supplemental Figure 2). These specific patterns in γH2AX levels were also seen using western blot analysis after nuclear fractionation (Figure 2D).

**Figure 2.**
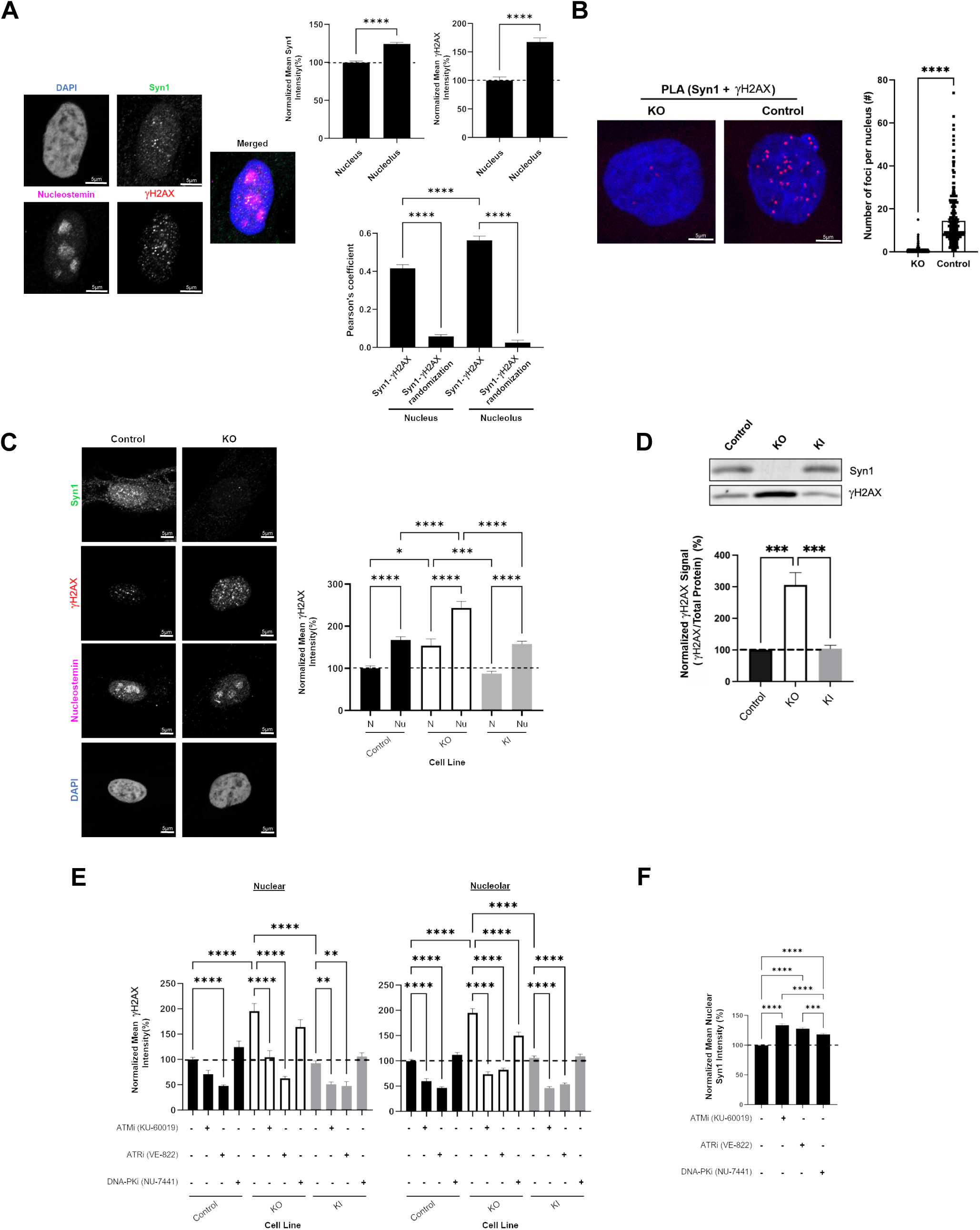
Alpha-synuclein interacts with γH2AX in the nucleolus and knocking out alpha-synuclein leads to increased ATM/ATR-driven H2AX phosphorylation. A, C) SK-Mel28 cells were seeded on PLL-coated coverslips and then fixed and stained for alpha-synuclein (Syn1), DSB marker (γH2AX), nucleolar mask (nucleostemin), and DAPI. Cells were imaged on the Zeiss 980 confocal microscope with airyscan and data was analyzed using FIJI (intensity) or Imaris (colocalization). Quantification represents quantification from 3 biological replicates. * p<0.05, *** p<0.001, **** p<0.0001 by T-test or ANOVA. Error bars denote SEM for all graphs. N=nucleus, Nu=nucleolus. Same γH2AX quantification of control cells between A and C. B) SK-Mel28 control and KO cells were seeded on PDL-coated coverslips and then fixed in 4% paraformaldehyde. Proximity Ligation Assay (Duolink) was completed using antibodies against Syn1 and γH2AX. Cells were imaged on the Zeiss 980 confocal microscope and number of foci per nucleus was measured using CellProfiler while masking for the nucleus using DAPI. Each figure shows representative images and quantification from 3 biological replicates. ****p<0.0001 by T-test. Error bars denote SEM. D) SK-Mel28 cells (control/KO/KI) were lysed and a nuclear fractionation was performed. Nuclear protein was run out on SDS-PAGE and probed for γH2AX and total protein. Western blots were imaged on Licor CLx imager. *** p<0.001 by ANOVA. Error bars denote SEM. Quantification from 4 biological replicates. E, F) SK-Mel28 cells were seeded on PLL-coated coverslips and treated with DMSO, KU-60019 (10µM), VE-822 (0.1µM), or NU-7441 (1µM) for 24 hours. Cells were fixed and stained for Syn1, γH2AX, nucleostemin and DAPI. Mean intensity of γH2AX signal within DAPI and nucleostemin masks analyzed using FIJI. ** p<0.01, *** p<0.001, **** p<0.0001 by ANOVA. Error bars denote SEM. Quantification from 3 biological replicates.

Three phosphoinositide 3-kinase (PI3K)-related kinases (PI3KK) family members are important for regulating γH2AX levels, ataxia-telangiectasia-mutated (ATM), ataxia-telangiectasia and Rad3 related (ATR), and DNA-dependent protein kinase (DNA-PK)^47^. Pharmacologic ATR inhibition reduced γH2AX levels in the nucleoplasm, and both ATM and ATR inhibition reduced γH2AX levels in the nucleolus in control SK-Mel28 cells, while DNA-PK inhibition had no effect in either compartment (Figure 2E). In αSyn KO cells, both ATM and ATR inhibition reduced γH2AX levels in the nucleus and nucleolus, while DNA-PK inhibition again had less prominent effects, and similar patterns were seen in αSyn KI-rescue cells to control cells (Figure 2E). Interestingly, inhibition of ATM, ATR, and DNA-PK all moderately increased nuclear αSyn levels (Figure 2F). Similar changes in nuclear γH2AX levels after ATM, ATR, and DNA-PK inhibition were also seen using the complementary In-Cell Western assay (Supplemental Figure 3C). Taken together, these results show that ATM and ATR are required for the increased phosphorylation of H2AX detected in the αSyn KO condition. Additionally, inhibiting any of these kinases – ATM, ATR, or DNA-PK –increases nuclear αSyn levels, potentially as part of a compensatory mechanism to improve DSB repair.

To directly test the role of αSyn in facilitating DSB repair, we induced DNA damage throughout the nucleoplasm and nucleolus (pan-nuclear) in SK-Mel28 cells with the chemotherapeutic agent bleomycin (100µg/ml, 1 hour). Bleomycin chemically induces DSBs, in addition to other forms of DNA damage like single-strand breaks^48^. As expected, bleomycin treatment increased pan-nuclear γH2AX immunocytochemical intensity (Figure 3A). Similar to our results with PI3KK inhibition, bleomycin also moderately increased pan-nuclear αSyn levels (Figure 3A). Pan-nuclear γH2AX levels were higher in αSyn KO cells compared to control cells treated with bleomycin and this was attenuated in αSyn KI-rescue cells (Figure 3A). Similar results were also seen using western blot (Figure 3B) and In-Cell Western (Figure 3C) analyses. Next, we used immunocytochemistry to compare γH2AX levels between the nucleoplasm and nucleolus after bleomycin treatment, as we previously did in untreated cells (Figure 2C). Bleomycin significantly increased γH2AX levels in both the nucleoplasm and in nucleoli (Figure 3E) with no clear differences between the two compartments, likely because bleomycin is a nonselective inducer of DNA damage throughout the nucleus with no preference for nucleolar versus nuclear DNA.

**Figure 3.**
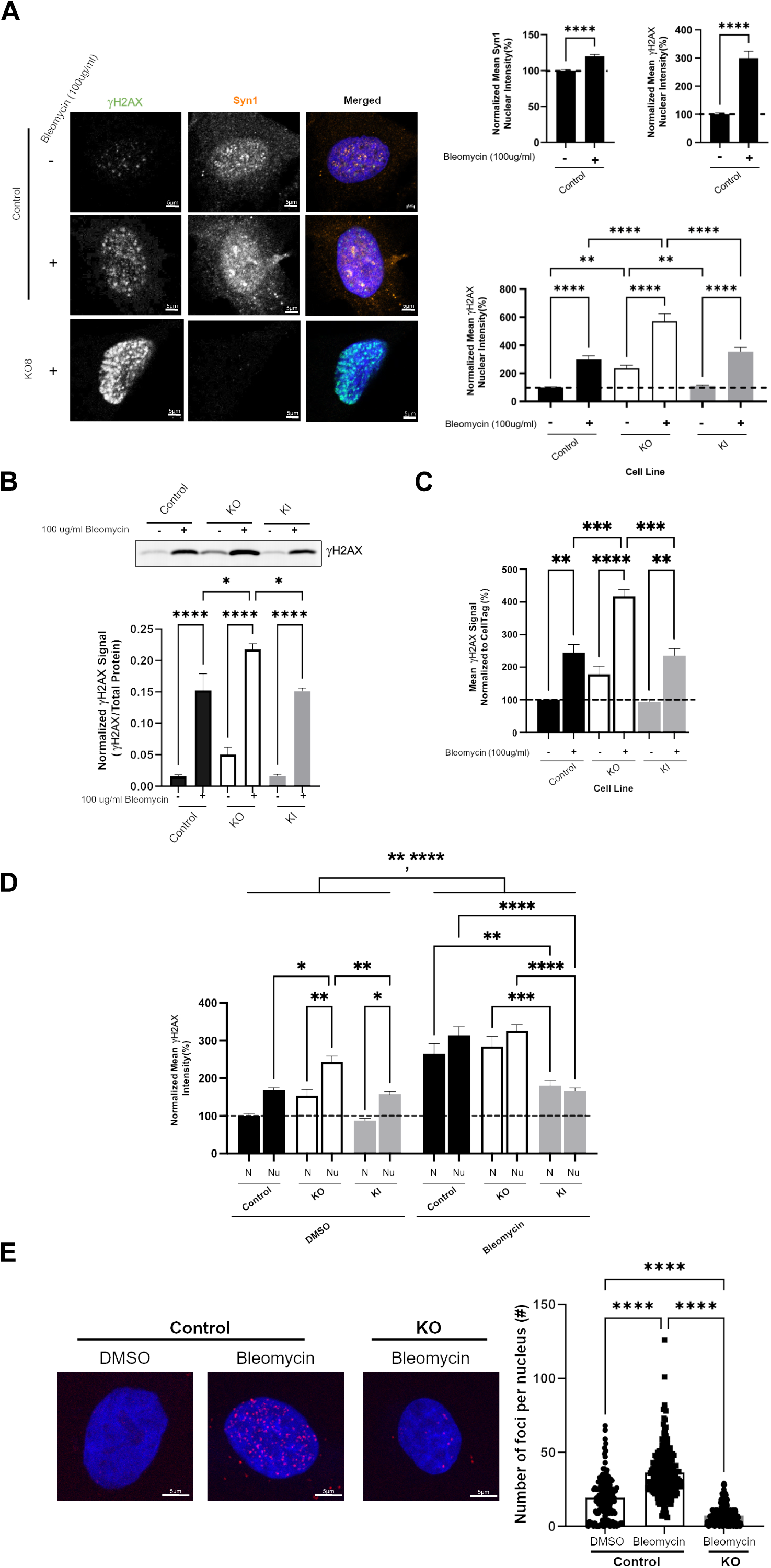
Alpha-synuclein is important in global DNA damage response pathways. A) SK-Mel28 cells (control/KO/KI) were seeded on PLL-coated coverslips and treated with DMSO or 100µg/ml bleomycin for 1 hour. Cells were fixed and stained for Syn1, γH2AX, and DAPI. Mean intensity of γH2AX signal within DAPI masks were analyzed using FIJI. ** p<0.01, **** p<0.0001 by ANOVA. Error bars denote SEM. Quantification from 3 biological replicates. Same γH2AX quantification between graphs with control cells. B) SK-Mel28 cells (control/KO/KI) were treated with DMSO or 100µg/ml bleomycin for 1 hour and lysed and a nuclear fractionation was performed. Nuclear protein was run out on SDS-PAGE and probed for γH2AX and total protein. Western blots were imaged on Licor CLx imager. * p<0.05, **** p<0.0001. Error bars denote SEM. Quantification from 4 biological replicates. C) SK-Mel28 cells were seeded in a black-welled PDL-coated 96 well plate and treated with DMSO or 100µg/ml bleomycin for 1 hour. Cells were processed according to the In-Cell Western manufacturer instructions and stained for γH2AX (800) and CellTag (700). Plates were imaged on the Licor CLx. ** p<0.01, *** p<0.001, **** p<0.0001 by ANOVA. Error bars denote SEM. Quantification from 3 biological replicates. D) SK-Mel28 cells (control/KO/KI) were seeded on PLL-coated coverslips and treated with DMSO or 100µg/ml bleomycin for 1 hour. Cells were fixed and stained for Syn1, γH2AX, nucleostemin, and DAPI. Mean intensity of γH2AX signal within nucleostemin and DAPI masks were analyzed using FIJI. * p<0.05, ** p<0.01, ***p<0.001, **** p<0.0001 by ANOVA. Error bars denote SEM. N=nucleus, Nu=nucleolus. Quantification from 3 biological replicates. E) SK-Mel28 cells were seeded on PDL-coated coverslips, treated with DMSO or 100µg/ml bleomycin for 1 hour, and fixed in 4% paraformaldehyde. DNA Damage In Situ Ligation Followed by Proximity Ligation Assay (Duolink) was completed as previously described (Galbiati and Fagagna, 2019) using a DNA linker and Syn1. Cells were imaged on the Zeiss 980 confocal microscope and number of foci per nucleus was measured using CellProfiler within nuclear masking with DAPI. Each figure shows representative images and quantification from 3 biological replicates. ****p<0.0001 by ANOVA. Error bars denote SEM.

In order to understand the spatial relationship between αSyn and DNA at the sites of DSBs, we used a recently developed modified PLA technique, DNA Damage In Situ Ligation Followed by Proximity Ligation Assay (DI-PLA)^49^. This approach detects a protein of interest, in our case αSyn, and a hairpin-shaped biotinylated DNA oligonucleotide that only ligates to DNA ends (found in DSBs) as the PLA partner. This allows sensitive detection of proteins located within ∼40nm of a DSB site. We found a higher number of αSyn DI-PLA foci in SK-Mel28 cells treated with bleomycin (Figure 3D), indicating that αSyn is located close to the sites of DSBs after bleomycin treatment. We next treated SK-Mel28 cells with hydroxyurea to test whether αSyn could protect against replication stress-induced DNA damage, as was previously shown to occur in yeast^50^. We did not find evidence that αSyn was important for repairing DNA damage induced by hydroxyurea (*data not shown*), as it is for bleomycin (Figure 3). Altogether, these data indicate that αSyn is upregulated in response to DNA damage and may play a role in modulating DSB repair and that this might be particularly important in the nucleolus.

### Alpha-synuclein regulates nucleolar double-strand break repair of ribosomal DNA

Given that bleomycin is a non-selective inducer of DNA damage both in the nucleoplasm and nucleolus, and our data suggests that αSyn is particularly enriched within the nucleolus, we next used an approach to create nucleolar-specific DSBs in rDNA with the intron-encoded endonuclease I-PpoI^51^, that recognizes a 13-15bp DNA sequence within the 28S rDNA coding region. I-PpoI-induced DSBs in rDNA leads to a large-scale reorganization of nucleolar structure, including the formation of nucleolar “caps” at the nucleolar periphery that allow DSB repair components that do not accumulate inside nucleoli to associate with damaged DNA for repair. After repair occurs, the nucleolus returns to its normal structure and caps disappear. After transfecting SK-Mel28 cells with mRNA for I-PpoI WT or the catalytically inactive mutant H98A (negative control), we measured a clear increase in nuclear γH2AX, with a pattern indicating nucleolar-specific DSBs in the I-PpoI WT condition (Figure 4A). There was also a significant increase in nuclear αSyn levels upon rDNA DSBs (Figure 4A), indicating that the cell may upregulate protein levels of αSyn as a consequence of rDNA damage, similar to that seen with bleomycin and PI3KK inhibition. Interestingly, γH2AX levels in the αSyn KO condition following induction of rDNA DSBs by I-PpoI were significantly increased compared to control or αSyn KI-rescue cells (Figure 4A). This increased γH2AX phenotype was rescued when αSyn was reintroduced into the KO background. A similar pattern was also seen using western blot analysis (Figure 4B). Although I-PpoI is used extensively to study rDNA damage, the recognition sequence for this endonuclease is found in a small number of genomic locations outside of rDNA. In order to further test the role of αSyn in rDNA DSB repair, we next used an alternative CRISPR/Cas9 system to induce DSBs at a site only found in rDNA using gRNAs targeting the 28S rDNA subunit^51, 52^. Similar to our I-PpoI experiments, CRISPR/Cas9-treated cells exhibited a high proportion of nucleolar caps (visualized by staining for the nucleolar protein Treacle) and increase in γH2AX signal directly surrounding the caps compared to a non-targeting gRNA negative control treated cells (Figure 4C). Similarly, γH2AX levels were increased in the αSyn KO condition, compared to control and αSyn KI-rescue cells, again indicating that αSyn loss-of-function impairs rDNA DSB repair (Figure 4C). To further test the role of nucleolar αSyn recruitment to sites of rDNA DSBs, DI-PLA analysis after nucleolar rDNA DSB induction with I-PpoI showed a higher number of αSyn DI-PLA foci compared to control conditions (Figure 4D), indicating that αSyn is recruited to within ∼40nm of rDNA DSBs. We also utilized laser-induced DNA damage using multiphoton laser illumination to generate small, sub-nucleolar regions of DNA DSBs using focused, short laser pulses, and then measured the immediate response of GFP-tagged human αSyn. Focal illumination of small nucleolar subregions with a short pulse of high intensity laser light induced the rapid redistribution of αSyn-GFP to the site of damage (Figure 4E).

**Figure 4.**
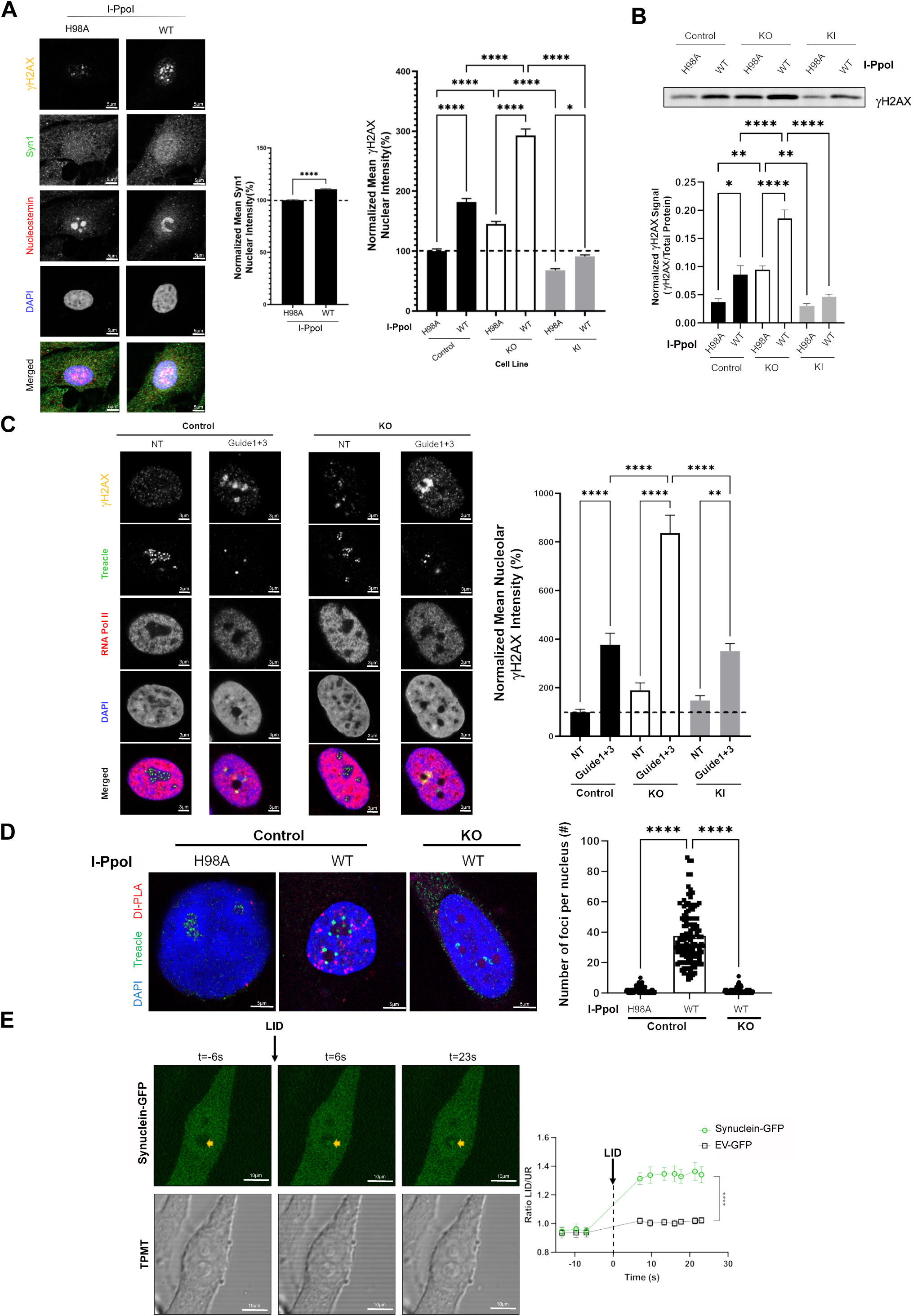
Inducing rDNA DSBs increases alpha-synuclein levels and localization to sites of damage and alpha-synuclein knockout significantly increases damage signature. A) SK-Mel28 cells (control/KO/KI) were seeded on PLL-coated coverslips and then transfected with WT and H98A I-PpoI mRNA. 6-hours post transfection, cells were fixed and stained for Syn1, γH2AX, nucleostemin, and DAPI. Cells were imaged on the Zeiss 980 confocal microscope and data was analyzed using FIJI. Quantification from 3 biological replicates. * p<0.05, ****p<0.0001 by ANOVA. Error bars denote SEM. B) SK-Mel28 cells (control/KO/KI) were transfected with WT and H98A I-PpoI mRNA. 6-hours post transfection, cells were lysed and a nuclear fractionation was performed. Nuclear protein was run out on SDS-PAGE and probed for γH2AX and total protein. Quantification from 4-6 biological replicates. *p<0.05, **p<0.01, ****p<0.0001 by ANOVA. Error bars denote SEM. C) SK-Mel28 cells (control/KO/KI) were seeded on PLL-coated coverslips and then transfected with Cas9 (TrueCut Cas9 Protein v2, Invitrogen) and guideRNAs that target portions of the 28S unit of rDNA or non-targeting control (NT vs. Guide 1+3). Twenty-four hours after transfection, cells were fixed and stained for γH2AX, treacle, RNA polymerase II, and DAPI. Cells were imaged on the Zeiss 980 confocal microscope and analyzed using RNA polymerase II anti-masking. γH2AX intensity was measured within a radius of treacle-identified nucleolar cap. Quantification from 3 biological replicates. ** p<0.01, ****p<0.0001 by ANOVA. Error bars denote SEM. D) SK-Mel28 cells (control/KO) were seeded on PDL-coated coverslips and then transfected with WT and H98A I-PpoI mRNA. 6-hours post transfection, cells were fixed and DNA Damage In Situ Ligation Followed by Proximity Ligation Assay (Duolink) was completed as previously described (Galbiati and Fagagna, 2019) using a DNA linker and Syn1. A treacle co-stain was included to identify nucleolar cap formation. Cells were imaged on the Zeiss 980 confocal microscope and number of foci per nucleus was measured using CellProfiler within nuclear masking with DAPI. Each figure shows representative images and quantification from 3 biological replicates. ****p<0.0001 by ANOVA. Error bars denote SEM. E) SK-Mel29 control cells expressing Synuclein-GFP or Empty Vector-GFP (EV). Yellow arrows show targeting of laser-induced damage (LID) pulse in the nucleolus. Baseline (t=-6s) and after LID (t=6 and 23s) images show accumulation of Synuclein-GFP at DNA damage site. Data from graph shows calculated enrichment ratio at LID site (compared to an adjacent site in the nucleolus). Quantification from >20 cells over 2 biological replicates. ****p<0.0001 by t-test. Error bars denote SEM.

We next investigated which PI3KK family member/s are responsible for the increase in γH2AX signal we detect in αSyn KO cells after I-PpoI induction of rDNA DSBs. This analysis revealed the largest decrease in γH2AX intensity in cells treated with ATM inhibitor, and a less pronounced, but still substantial, decrease after ATR or DNA-PK inhibition (Figure 5A). These results differ from baseline (Figure 2E), in that ATR-inhibition no longer had as drastic of an effect on γH2AX signal attenuation. This could be due to the more dominant incidence of replication-stress induced DSBs during homeostatic conditions that are controlled by ATR cascades, as opposed to a shift towards ATM cascades during I-PpoI-induced DSBs. These results indicate that the increase in γH2AX phosphorylation in control SK-Mel28 cells after I-PpoI WT treatment is driven primarily by ATM activity.

**Figure 5.**
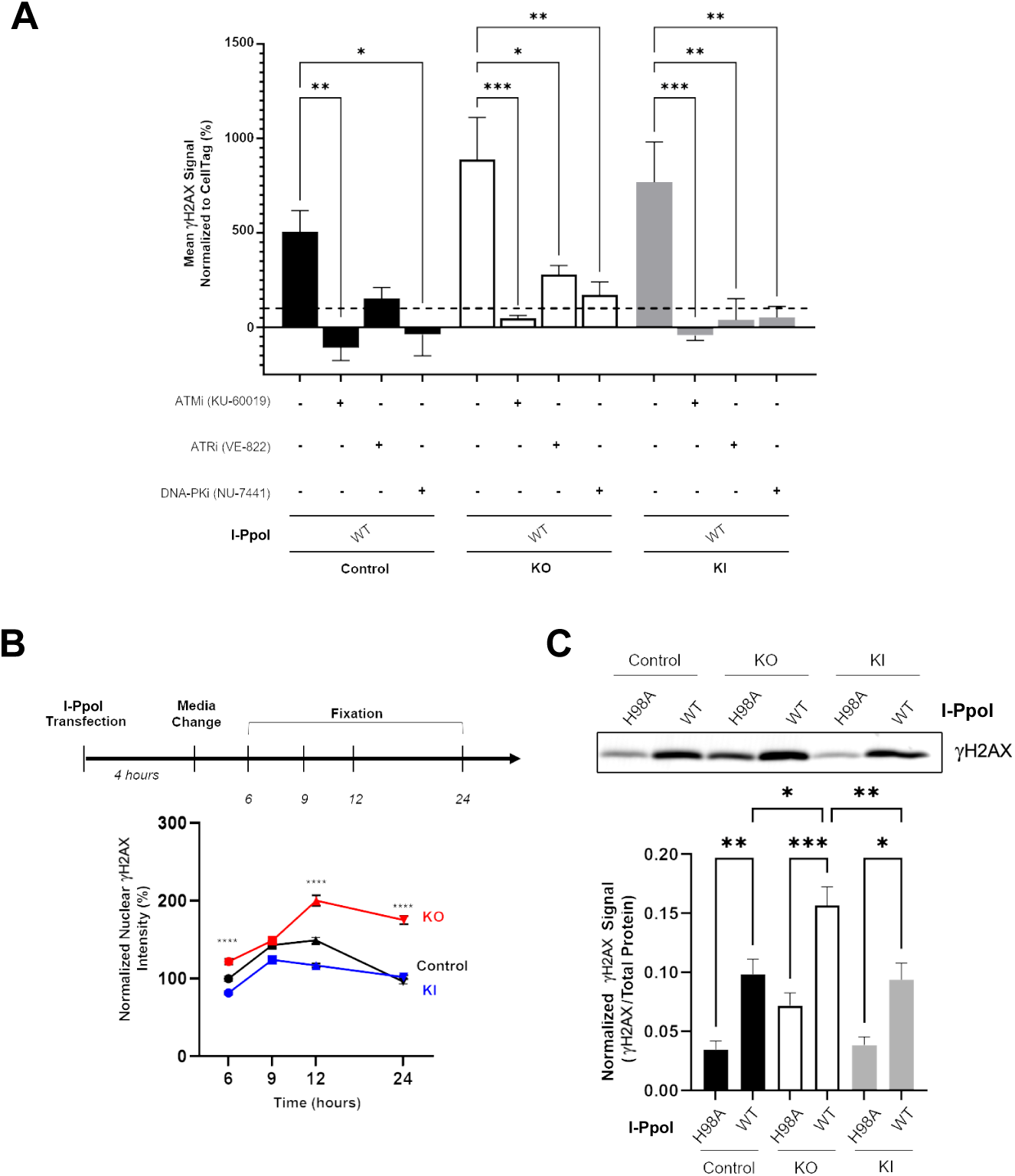
Alpha-synuclein knockout significantly impairs recovery of rDNA damage in an ATM-dependent manner. A) SK-Mel28 cells were seeded in a black-welled PDL-coated 96 well plate and treated with DMSO, KU-60019 (10µM), VE-822 (0.1µM), or NU-7441 (1µM) for 24 hours. Cells were then treated with WT and H98A I-PpoI mRNA for 6 hours in the presence of the inhibitors. Cells were processed according to the In-Cell Western manufacturer instructions and stained for γH2AX (800) and CellTag (700). Plates were imaged on the Licor CLx. * p<0.05, ** p<0.01, *** p<0.001 by ANOVA. Error bars denote SEM. Quantification from 6 biological replicates. Normalization to control cells transfected with I-PpoI H98A and treated with DMSO. B) SK-Mel28 cells (control/KO/KI) were seeded on PLL-coated coverslips and then transfected with WT and H98A I-PpoI mRNA. At the indicated time point post transfection, cells were fixed and stained for Syn1, γH2AX, nucleostemin, and DAPI. Cells were imaged on the Zeiss 980 confocal microscope and data was analyzed using FIJI. Quantification from 3 biological replicates. Statistical labeling denotes significance between control and KO cell lines, ****p<0.0001 by ANOVA. C) SK-Mel28 cells (control/KO/KI) were transfected with WT and H98A I-PpoI mRNA. After 24 hours, cells were lysed and a nuclear fractionation was performed. Nuclear protein was run out on SDS-PAGE and probed for γH2AX and total protein. Quantification from 5 biological replicates. *p<0.05, **p<0.01, ***p<0.001, by ANOVA. Error bars denote SEM.

To understand the potential importance of αSyn in the kinetics of repair after I-PpoI-induced rDNA DSB formation, we measured the time course of γH2AX changes. Using immunocytochemistry, we found a delay in recovery of γH2AX levels after I-PpoI transfection in αSyn KO cells that lasts at least 24 hours compared to control or αSyn KI-rescue cells (Figure 5B). Western blot analysis of γH2AX at the 24-hour time point also confirmed a persistent elevation of γH2AX in αSyn KO cells compared to either control group (Figure 5C). These data strongly suggest that αSyn is important in regulating the DDR in a way that facilitates DSB repair and that its loss-of-function leads to higher levels of DNA damage that cells are slower to repair.

### Alpha-synuclein is recruited near nucleolar caps and regulates the rate of DSB repair

I-PpoI-treated SK-Mel28 cells were studied to understand the spatial distribution of αSyn after rDNA DSB induction and nucleolar cap formation. Discrete αSyn foci were found directly adjacent to the nucleolar cap (Figure 6A), similar to previous reports for 53BP1 and BRCA1 after rDNA DSB formation^51, 53, 54^, with the most αSyn signal within 2 microns of the nucleolar cap. Actinomycin-D-mediated RNA Pol I inhibition, which also induces nucleolar caps by a mechanism independent of DNA damage, did not show the same αSyn localization to the juxta-nucleolar cap region (Figure 6A). To better understand how αSyn regulates DSB repair and nucleolar cap kinetics, we next utilized longitudinal live-cell imaging techniques. GFP-tagged treacle-expressing control, αSyn KO, and αSyn KI-rescue cells were transfected with I-PpoI mRNA to induce rDNA DSBs and nucleolar dynamics were visualized over a ∼12-hour period (∼5-18 hours after I-PpoI transfection). No significant differences in percentage of cells with nucleolar caps or the time to cap formation were detected (Figure 6B). Interestingly, however, the rate at which nucleolar caps reorganized back to normal nucleoli was slower in αSyn KO cells compared to control or αSyn KI-rescue cells (Figure 6B). Similar to our γH2AX time course experiments (Figure 5A-B), this live-cell imaging data also strongly suggests that αSyn is important in regulating the rate at which DSBs in rDNA are repaired.

**Figure 6.**
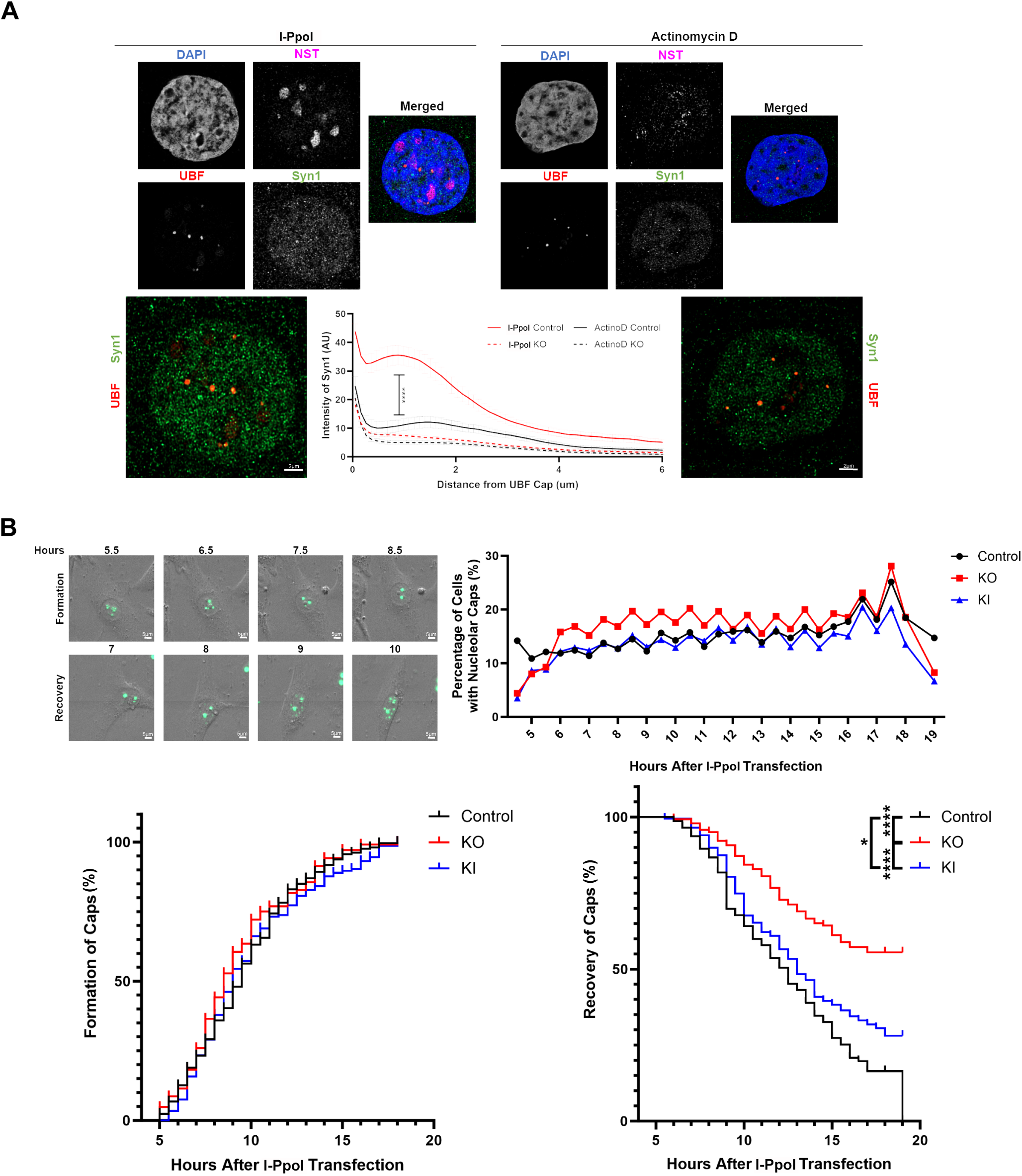
Alpha-synuclein localizes to DSB-induced nucleolar caps and is important for nucleolar cap recovery. A) SK-Mel28 cells were seeded on PLL-coated coverslips and then treated with WT I-PpoI mRNA for 6 hours or 100ng/ml Actinomycin D for 1 hour prior to fixation. Cells were stained for Syn1, nucleolar caps marker (UBF), nucleostemin, and DAPI. Cells were imaged on the Zeiss 980 confocal microscope with Airyscan oversampling and Joint Deconvolution and Channel Alignment post-processing. 3D renderings and distance from cap analysis were produced using Imaris software. Quantification from 5 biological replicates. Error bars denote SEM. Statistical significance was calculated via nonlinear regression (99% confidence interval). B) SK-Mel28 cells (control/KO/KI) were seeded on PDL-coated 8-well Ibidi plates. Cells were transfected with 800ng GFP-Treacle using Lipofectamine 3000. Twenty-four hours post-transfection, cells were treated with WT I-PpoI mRNA. Four hours after treatment, live-cell imaging was performed using the Zeiss Celldiscoverer 7 and imaged for 15 hours. Quantification from 5 biological replicates. *p<0.05, ****p<0.0001 by two-way ANOVA or Mantel-Cox test.

### Alpha-synuclein knockout impairs 53BP1 recruitment to nucleolar caps and leads to micronuclei formation and growth dysregulation

Given our data suggesting that γH2AX levels are persistently elevated after DSB formation in the αSyn loss-of-function condition (Figures 3-5), we next tested whether this could be due to a deficiency in a step downstream of γH2AX during DDR signaling. 53BP1 has been previously shown to be recruited near nucleolar caps after specific rDNA DSBs induction^51, 53, 54^ and downstream of H2AX phosphorylation. 53BP1 is important for DSB repair pathway choice, generally promoting non-homologous end joining (NHEJ) and limiting homologous recombination (HR)^55^. However, DSB repair in heterochromatin by HR also requires 53BP1 suggesting that the role of 53BP1 may vary dependent on the context^56^. We found a decrease in 53BP1 near the nucleolar cap in αSyn KO cells compared to control or αSyn KI-rescue cells after I-PpoI transfection (Figure 7A), suggesting that αSyn is important for 53BP1 recruitment to DSBs downstream of γH2AX in nucleolar DDR signaling. We next tested for possible downstream cellular consequences of delayed DSB repair in αSyn KO cells. Importantly, there are previous investigations suggesting that progressing through mitosis with damaged rDNA leads to abnormal nuclear morphology in cells, like micronuclei^52^, and that 53BP1 dysregulation may influence micronuclei formation^57, 58^. We measured an increase in the percentage of cells with micronuclei at baseline in the αSyn KO condition, that was also present 6- and 24-hours post rDNA DSB induction with I-PpoI (Figure 7B). We also measured proliferation, migration, and invasion to determine downstream contributions of αSyn function to these cellular phenotypes, since previous studies have found dysregulation of various growth measurements in αSyn KO cells^40, 59^. SK-Mel28 αSyn KO cells also exhibited impaired proliferation, migration, and invasion capabilities compared to control and αSyn KI-rescue cells (Figure 7C), suggesting that αSyn-mediated rDNA DSB repair is important for cell survival and growth.

**Figure 7.**
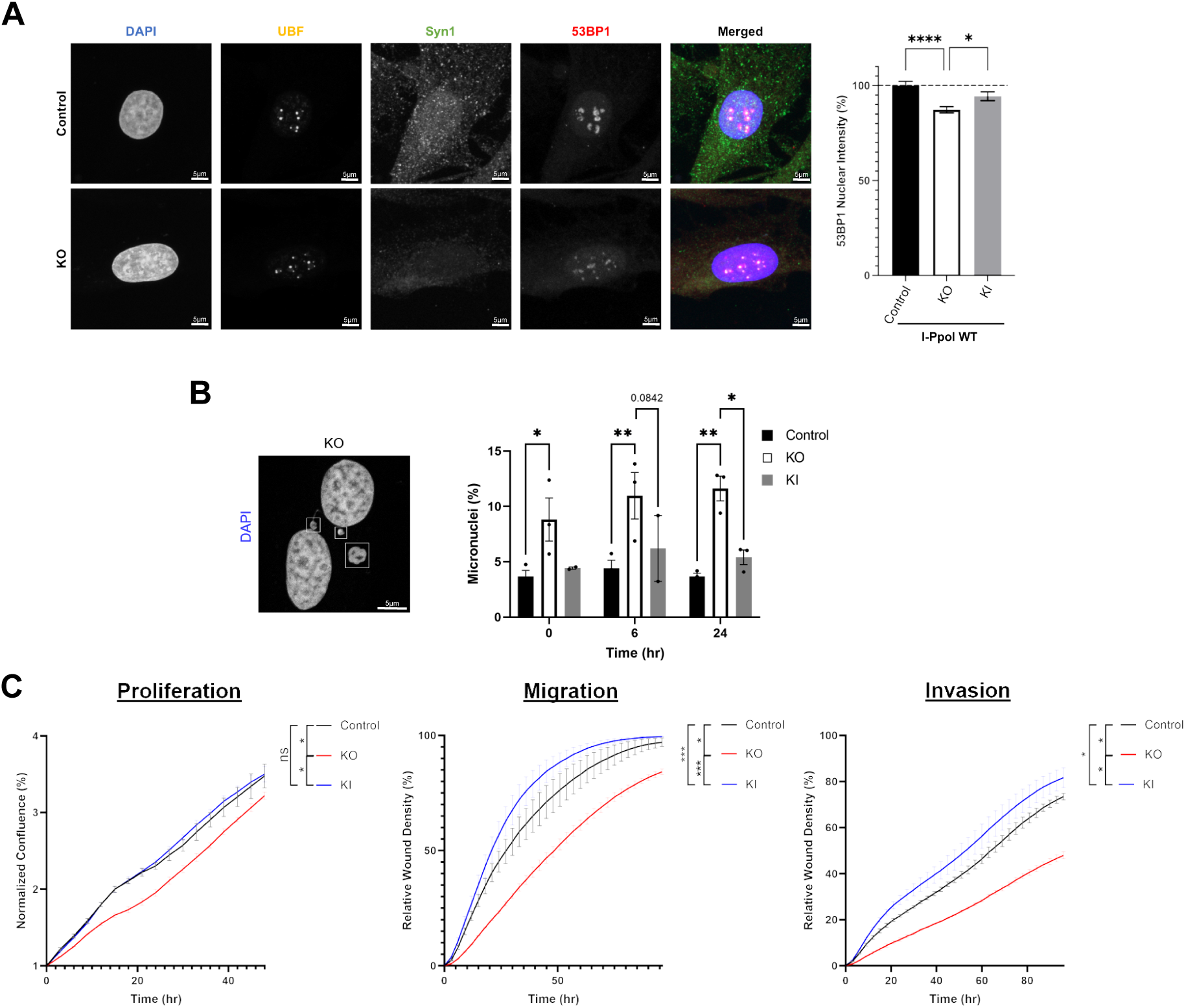
Alpha-synuclein knockout leads to dysregulated recruitment of 53BP1 to nucleolar caps and significantly increases occurrences of micronuclei formation with impaired cellular growth phenotypes. A) SK-Mel28 cells (control/KO/KI) were seeded on PLL-coated coverslips and then transfected with WT I-PpoI mRNA. 6-hours post transfection, cells were fixed and stained for Syn1, UBF, 53BP1, and DAPI. Cells were imaged on the Zeiss 980 confocal microscope and data was analyzed using FIJI. Quantification from 4 biological replicates. * p<0.05, ****p<0.0001 by ANOVA. Error bars denote SEM. B) SK-Mel28 cells were seeded on PLL-coated coverslips and then transfected with WT I-PpoI mRNA. 0, 6, and 24-hours post transfection, cells were fixed and stained for DAPI. Cells were imaged on the Zeiss 980 confocal microscope. Micronuclei were hand counted with the experimenter blinded to cell condition. Quantification from 3 biological replicates. *p<0.05, **p<0.01 by ANOVA. Error bars denote SEM. C) SK-Mel28 cells were plated at 6,000 cells per well on 80μg/ml Matrigel coated ImageLock plates. Graphs represent temporal progression of total confluence (proliferation), or wound closure using Relative Wound Density as the metric to measure migration or invasion. Data represents time-course of means of each cell line among 3 biological replicates. *95%CI, ***99%CI by nonlinear regression (migration) or simple linear regression (proliferation and invasion) analysis.

### Transcriptomic analysis reveals alpha-synuclein involvement in nucleolar DNA binding regulation and damage repair pathways

Previous work has shown that rDNA DSB induction leads to transcriptional silencing of rDNA via an ATM-dependent process important for nucleolar cap formation^52, 53, 60-63^, and other work has implicated αSyn in transcriptional regulation, both in the nucleolus^64, 65^ and in the context of neurodegeneration^66^. Therefore, we next tested for a potential role of αSyn in rDNA transcriptional silencing after DSB induction. Measurements of transcription using 5-EU (5-Ethynyl Uridine) incorporation in SK-Mel28 cells treated with gRNAs targeting rDNA or Actinomycin D (Act D) were compared between control, αSyn KO, and rescue (KI) backgrounds. Our results suggest that αSyn plays no detectible role in the specific silencing of rDNA transcription that occurs after DSB induction, but that it may have effects on general rDNA transcription, since this was increased in the αSyn KO condition at baseline (Supplemental Figure 4A, B).

In order to better understand these possible changes in baseline rDNA transcription in the αSyn KO cells and what cellular processes might be altered, whole-cell RNA extraction and bulk sequencing (RNAseq) analysis was performed in SK-Mel28 control and αSyn KO cells. Differentially expressed transcripts were identified and assigned p-values and false discovery rates (FDR)^67^. A gene set was selected that included all immunofluorescence-validated nucleolar genes^68^ and cross-referenced to our RNAseq differential gene expression list. 64 nucleolar-associated genes were identified to be significantly up- or down-regulated in the αSyn KO line compared to the control cells (Table 1). We identified 37 upregulated (log_2_-fold change 2.01 to 29.53) and 27 downregulated (log_2_-fold change -2.00 to -8.07, Figure 8A) nucleolar-associated gene transcripts. Six of the transcripts exhibiting the largest changes were validated by qRT-PCR and all showed changes in the expected direction (Figure 8A, Table 2), with *ATF3*, *DTX3*, and *HMGA2* significantly upregulated and *CRIP2* significantly downregulated (Supplemental Figure 4C). *ATF3* (Activating Transcription Factor 3), which showed the largest upregulation in αSyn KO cells both by RNAseq and qRT-PCR, has been previously implicated in cellular responses to a variety of stresses^69^ and to be important for regulating DSB repair^70, 71^. Further gene ontology analysis revealed upregulated transcripts associated with DNA binding (*HMGA1, HMGA2*) and transcription (*SFRP1, HMGA1, HMGA2, EN1, TBL1X, BMP7, ATF3)* and downregulated gene populations associated with RNA polymerase DNA binding (*ZNF397, ZNF419, FOXJ2, ZBTB43, ZNF689, ZNF33B, STOX1*) (Figure 8B).

**Figure 8.**
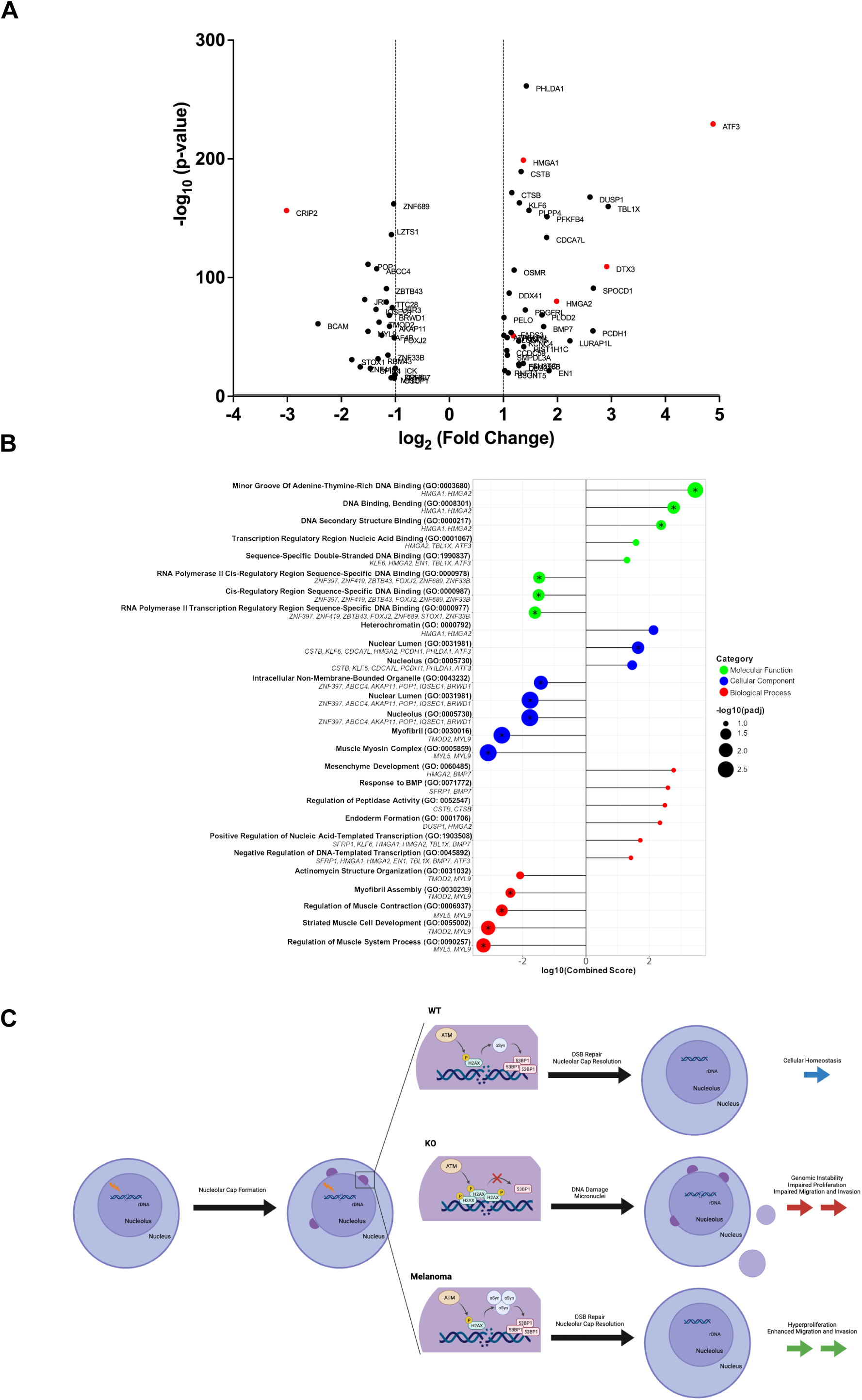
Transcriptomic analysis reveals alpha-synuclein involvement in nucleolar DNA damage pathways. A) Total RNA was extracted from SK-Mel28 control and KO cells and sent for RNA-sequencing analysis at Indiana University. Differentially expressed gene transcripts in KO cells compared to control were identified. These were cross-referenced to over 500 nucleolar-specific genes. 64 genes were identified and plotted on a volcano plot using p-value and fold change. All transcripts, greater than 1 standard deviation and less than -1 standard deviation were plotted. Red labelling indicated RT-PCR validated genes. B) All significant upregulated and downregulated transcripts underwent gene ontology analysis, which included pathways associated with molecular function, cellular component, and biological process. Gene sets found to be significant after Benjamini-Hochberg multiple testing correction (P.adj < 0.05) were marked with an asterisk. C) Graphic outlining mechanistic function of alpha-synuclein in the nucleolar DNA damage response pathway.

## Discussion

In this study, we have expanded our knowledge of the nuclear functions of the neurodegeneration and cancer-associated protein, αSyn, by showing its particular importance in facilitating nucleolar DDR pathways in melanoma. Our data suggests that αSyn is preferentially enriched within the nucleolar granular component, where it colocalized with the marker of DSB repair γH2AX. αSyn loss-of-function by genetic deletion increased γH2AX levels in the nucleolus via an ATR- and ATM-dependent pathway. Pan-nuclear DNA damage induction via bleomycin treatment increased global nuclear αSyn, while αSyn KO exacerbated the increase in γH2AX throughout the nucleolus and nucleoplasm. Selective rDNA DSB induction within the nucleolus via I-PpoI- or CRISPR/Cas9-based approaches both increased nuclear αSyn and γH2AX, and in αSyn KO cells this γH2AX increase was even greater and mediated by ATM. Our DI-PLA analysis showed that αSyn is present within close proximity (∼40nm) of these rDNA DSBs. Kinetic studies of the γH2AX response after selective rDNA DSB induction also revealed a significant delay in γH2AX resolution in αSyn KO cells, which was also associated with impaired resolution of nucleolar caps. αSyn KO led to a reduction in 53BP1 recruitment to the nucleolar caps after inducing rDNA DSBs. These abnormalities in nucleolar DSB repair were associated with multiple downstream cellular effects, including increased micronuclei formation, impaired proliferation, migration, and invasion. Lastly, transcriptomic analysis of αSyn KO cells reinforced our finding that αSyn loss-of-function leads to dysregulated DSB repair. Altogether, these findings illuminate a new role for αSyn in the nucleolar DDR where it acts downstream of ATM-mediated H2AX phosphorylation and is important for facilitating 53BP1 recruitment. αSyn loss-of-function leads to delayed DSB repair with specific cellular consequences (Figure 8C).

Although our findings strongly suggest that αSyn acts downstream of H2AX phosphorylation to enable proper 53BP1 recruitment to DSBs, the specific mechanism by which αSyn helps mediate this recruitment remains unknown. Previous work has shown that 53BP1 recruitment to damaged sites requires H2AX K13-K15 ubiquitination by the RING-type E3 ubiquitin ligases RNF8 and RNF168^72-75^, and histone H4 K20 methylation^76, 77^. Interestingly, expression of the PD-associated A53T αSyn point mutation has also been recently shown to cause delayed repair and abnormal RNF8 retention at DSB repair foci after ionizing radiation treatment^78^, which is similar to our results with αSyn KO cells. Protein methylation is another post-translation modification important for 53BP1 recruitment during the DDR and very recent work identified PRMT5 (Protein Arginine Methyltransferase 5) as an αSyn interactor through proteomics analysis^79^. PRMT5 can directly methylate 53BP1 in its GAR domain to facilitate docking of 53BP1 to DSB sites^80, 81^. It will be interesting to test in future studies whether αSyn regulates the histone post-translational modification cascade mediated by RNF8/168, PRMT5 mediated 53BP1 methylation, and/or other pathways to directly facilitate 53BP1 binding to DSBs.

Our RNA-seq analysis identified 64 nucleolar-associated transcripts that were up or down regulated in αSyn KO cells, many of which have direct links to the DDR. Gene ontology analysis also showed enrichment of DNA binding and transcriptional regulation pathways (Figure 8B). ATF3, which had the largest change of any identified transcript (∼30-fold increase in αSyn KO cells) is particularly interesting, since it is activated upon DNA damage in a p53-dependent manner and its overexpression moderately suppresses cell growth^82-84^ and it facilitates DSB repair^70, 71^. HMGA1 and HMGA2 are small non-histone proteins that can bind to DNA and modify chromatin state and were also upregulated in our dataset 2.5- and 4-fold, respectively. Both have been implicated in the DDR and are direct ATM/ATR kinase targets^85-88^. HMGA2 is associated with suppression of NHEJ via hyperphosphorylation of DNA-PKcs^89^, and interestingly, our data suggests αSyn KO reduces 53BP1 recruitment. These RNA-seq data are consistent with the concept that αSyn is important in melanoma cells for determining the mechanism of DSB repair, which will also be important to test further in future studies.

Our data fits into a larger landscape of links between the nucleolus, genomic instability, and cancer. Due to their highly proliferative nature, cancers are vulnerable to replicative exhaustion and subsequent genome instability. The nucleolus is especially prone to this due to its extremely high levels of transcriptional activity of repetitive rDNA sequences. Estimates suggest that up to 60% of all transcription within a normal mammalian cell occurs at rDNA^90^, and faithful ribosome biogenesis is even more important in malignant cells to sustain their increased levels of cell division and growth^91, 92^. In melanoma specifically, substantial evidence suggests that there is upregulation of DSB repair pathways to promote increased DSB repair capacity critical for survival^93, 94^. Evidence for this includes melanoma’s relative resistance to ionizing radiation^95, 96^ and that DSB repair inhibitors are particularly effective in many treatment models^97, 98^. Our findings that αSyn modulates nucleolar DSB repair and its loss-of-function negatively impacts cellular growth suggest that upregulation of αSyn levels in melanoma may also be part of a similar mechanism to improve DSB repair, allowing these cells to evade the programmed cell death and senescence pathways that would normally be triggered by high DSB levels.

It is well-established that PD patients and their first-degree relatives are at increased risk of melanoma, and symmetrically that melanoma patients are at increased risk of PD. Our previous work suggests that genetic or environmental factors that cause increased αSyn expression within certain individuals would predispose their post-mitotic neurons to accumulate cytoplasmic Lewy pathology and this, counterintuitively, triggers a loss of soluble, functional αSyn from the nucleus. This could lead to deficient DSB repair that contributes to programmed cell death^35, 36^. Our current data suggest these same individuals could also be predisposed to develop melanoma via a gain-of-function mechanism where increased αSyn levels improve DSB repair capacity within the nucleolus, which limits the senescence and programmed cell death pathways that are triggered by excessive DSBs associated with oncogenesis. This provides a novel framework for understanding the link between PD and melanoma and offers new potential therapeutic targets in melanoma that are focused on reducing αSyn-mediated nucleolar DSB repair.

## Methods

### Cell lines

The SK-Mel28 cell line were gifted by Dr. Stephan Witt (Louisiana State University), after being purchased from ATCC and authenticated at the University of Arizona Genetics Core via their STR Profiling Cell Authentication service. Per Shekoohi et al. 2021, αSyn knockout cells were created through CRISPR/Cas9 genome editing targeting *SNCA^40^*. In addition, re-expression of αSyn in the *SNCA* KO clone was established using lentivirus transduction of human αSyn under the CMV promoter. All SK-Mel28, A375, and PIG1 cell lines were cultured in appropriate medium suggested by ATCC. Primary melanocytes were isolated from patient foreskin samples provided by Oregon Health and Science University. Isolation protocol followed the steps outlined in^99^. All cells were maintained in a humidified chamber with constant supply of 5% CO_2_ and 95% O_2_ at 37C.

### Immunocytochemistry staining

SK-Mel28 cells were seeded onto poly-l-lysine treated glass coverslips and treated as indicated. Cells were then washed with 1x PBS and fixed using 4% paraformaldehyde for 15 minutes. After one wash in 1x PBS, cells were permeabilized in 0.25% Triton X-100 in PBS for 5 minutes. Coverslips were blocked in 10% goat serum/0.1% Triton X-100 in PBS for 30 minutes and then placed in the primary antibody overnight at RT. The next morning, cells were washed three times in 1x PBS and placed in secondary antibody overnight at RT. The following day, coverslips were washed 4 times in 1x PBS. The third wash contained DAPI (2.5µg/ml) for 20min. Coverslips were mounted using CFM2 antifade reagent and sealed with BioGrip. All immunofluorescence images were taken on a Zeiss Laser-Scanning Confocal Microscope 980 with Airyscan and analyzed with FIJI (2D analysis) or Imaris (3D analysis). Mean intensity was measured after imposing DAPI, RNA Polymerase II, or Nucleostemin masks over each cell. All cells within a 63x image was analyzed (∼3000cells/condition/biological replicate). Statistical significance was assigned using one-way ANOVA with multiple comparisons.

Ultra-resolution imaging samples were processed as described above and imaged on Zeiss Laser-Scanning Confocal Microscope 980 with Airyscan oversampling parameters. These images underwent Airyscan Joint Deconvolution and Channel Alignment post-processing steps, prior to Imaris 3D analysis.

Antibody specifics were as follows: Syn1 (BD Biosciences #610786, 1:500), Nucleophosmin (Abcam #52644, 1:100), Nucleostemin (Santa Cruz #166430, 1:500), Treacle (Millipore Sigma #HPA038237, 1:200), Fibrillarin (Abcam #5821, 1:100), RPA194 (Santa Cruz #48385, 1:200), γH2AX (Cell Signaling #9718, 1:500), Treacle (Sigma-Aldrich #GW22821, 1:10,000), RNA Pol II (Santa Cruz #47701, 1: 500), UBF (Millipore Sigma #HPA006385, 1:10,000), 53BP1 (BD Biosciences #612522, 1:1000).

### DNA Damage In Situ Ligation followed Proximity Ligation Assay (DI-PLA)

This protocol is adapted from Galbiati et al^49^. SK-Mel28 sells were grown on 13mm poly-d-lysine treated coverslips and fixed in 4% PFA for 15 minutes at room temperature followed by two washes with 1x PBS.

#### DI-PLA: Blunting

Coverslips were washed twice for 5 minutes with NEB2 buffer (50mM NaCl, 10mM Tris-HCl pH 8, 10mM MgCl_2_, 1mM DTT, 0.1% Triton X-100) and twice for 5 minutes with Blunting buffer (50mM NaCl, 10mM Tris-HCl pH 7.5, 10mM MgCl_2_, 5mM DTT, 0.025% Triton X-100). Coverslips were then inverted onto a 35μL drop on parafilm of NEB Blunting Reaction (NEB, E1201): (1mM dNTPs, 1X Blunting Buffer, 0.2mg/mL BSA, 1X Blunting Enzyme). Coverslips were incubated in a dark humidity chamber for 1hr at room temperature.

#### DI-PLA: Ligation

Coverslips were washed twice for 5 minutes with NEB2 buffer (50mM NaCl, 10mM Tris-HCl pH 8, 10mM MgCl_2_, 1mM DTT, 0.1% Triton X-100), then twice for 5 minutes with ligation buffer (50mM Tris-HCl pH 7.5, 10mM MgCl_2_, 10mM DTT, 1mM ATP). Coverslips were then inverted onto a 50μL drop on parafilm of Ligation Reaction (0.1μM DI-PLA Linker, 1X T4 Ligation Buffer (NEB, B0202), 1mM ATP, 0.2 mg/mL BSA, 1X T4 Ligase (NEB, M0202)) overnight at 4°C in dark humidity chamber followed by proximity ligation assay between biotin and protein of interest.

DI-PLA Linker: 5’-TACTACCTCGAGAGTTACGCTAGGGATAACAGGGTAATATAGTTT [BtndT] TTTCTATATTACCCTGTTATCCCTAGCGTAACTCTCGAGGTAGTA -3’.

#### Proximity Ligation Assay

Proximity Ligation Assay was performed without deviation from manufacturer’s instructions (DUO92008). Coverslips were washed in a 0.5mL volume and reactions were performed by inverting the coverslip onto a 35μL drop on parafilm. Following the proximity ligation reaction, cells were stained with DAPI (0.2µg/mL) for 3 minutes followed by one wash in PBS and one water wash. The cells were then inverted and mounted on glass coverslips with 15μL of prolong gold mounting media (LifeTech, P36934) & were cured overnight in the dark at room temperature.

All immunofluorescence images were taken on a Zeiss Laser-Scanning Confocal Microscope 980 and analyzed with CellProfiler. All cells within a 63x image was analyzed (∼30cells/condition/biological replicate). Statistical significance was assigned using t-test or one-way ANOVA with multiple comparisons.

### Transmission electron microscopy

SK-Mel28 control and KO cell pellets were fixed with 0.1M sodium cacodylate buffer (pH 7.2) containing 0.05% glutaraldehyde, 4% paraformaldehyde, and 0.1% picric acid for 2 hours at RT. The pellet was then processed for immuno-gold electron microscopy, using a microwave tissue processor (Pelco Biowave, Ted Pella, Inc, Redding, CA) as previously reported^100^. The pellet was gently removed from the tube and transferred to specimen dishes (Ted Pella, Inc). Briefly, the pellet was exposed to 1% osmium tetroxide/1.5% potassium ferricyanide in the Biowave (see Moore *et al*., 2020, for details), washed in water, then followed by 0.5% aqueous uranyl acetate, dehydrated in alcohol/propylene oxide, and embedded in Epon/Spurr resin. The pellet was thin sectioned (60 nm) on an ultramicrotome (Leica EM UC7, Buffalo Grove, Il), using a diamond knife (Diatome, Hatfield, PA). The sections were placed on 75 mesh grids, then incubated overnight using an antibody against alpha-synuclein (abcam, Boston, MA, #AB138501, rabbit polyclonal, 1:75) in TBST (tris buffered saline triton, pH 7.6) blocking solution (0.05% normal goat serum). The sections were then incubated in a secondary antibody (goat anti-rabbit, 1:50, in TBST 8.2, Jackson ImmunoResearch, West Grove, PA) tagged with a 12 nm gold particle for 90 minutes at room temperature. The sections were then viewed on a JEOL electron microscope (1400 TEM, JEOL, Peabody, MA). Photographs (digital camera, AMT, Danvers, MA) were taken of immuno-gold labeling of the nucleolus. The density of gold labeling was quantified as #/µm^2^ of nucleolar area.

### Western blot analysis

SK-Mel28 cells were seeded on 10cm plates to be ∼80% confluent the day of treatment. Cells were treated with bleomycin (100µg/ml) for 1hour or I-PpoI WT or H98A mRNA (7µg) as detailed below. After treatment, media was removed and cells were washed 1x with ice cold PBS. Cells were harvested by trypsinization, collected into 15ml conical tubes, and pelleted for 5min 200rfc. Liquid was aspirated, pellets were resuspended in 2ml PBS and transferred to 2ml microcentrifuge tubes. Proteins were extracted into cytosolic and nuclear fraction using the NE-PER extraction kit (Thermo-Fisher, cat 78833) according to the manufacturer’s recommendations with the addition of a brief sonfication (10 seconds, 10 kHz) after the first nuclear resuspension step. Protein preps were stored at -80C until Western blot analysis. 10-30µg protein was run on a 10-20% Tris-Glycine 1.0 mm gradient gel (Invitrogen) and transferred onto a Biodyne B Nylon Membrane (Thermofisher Scientific) at 30V for 2 hours on ice in 0.5% TBE using the Novex XCell II Blotting System (Invitrogen). If completing Syn1 staining, membranes are fixed in 4% paraformaldehyde/0.01% glutaraldehyde in PBS for 10 minutes at RT. Membranes were blocked overnight in Odyssey PBS Blocking Buffer (Li-Cor) and stained for 2 hours at RT with Syn1 (1:1,000; Biolegend) or γH2AX (1:1,000; Cell Signaling) and 1 hour at RT with IRDye 680CW Goat anti-mouse (1:10,000; Li-Cor) or IRDye 800CW Goat anti-rabbit (1:5,000; Li-Cor). All staining was normalized to total protein (Revert 700 Total Protein Stain, Licor). Images were acquired using Li-Cor Odyssey CLx Imaging System.

### In-cell western

SK-Mel28 cells were seeded on poly-d-lysine treated 96-well plates to be ∼80% confluent the day of treatment. Cells were treated as indicated. After treatment, media was removed and cells were fixed using 4% paraformaldehyde for 15 minutes. After one wash in 1x PBS, cells were permeabilized in 0.25% Triton X-100 in PBS for 5 minutes and blocked in 10% goat serum/0.1% Triton X-100 in PBS for 30 minutes and then placed in the primary antibody overnight at RT (γH2AX, Cell Signaling, 1:500). The next morning, cells were washed three times in 1x PBS and placed in secondary antibody at RT for 2 hours (IRDye 800CW Goat Anti-Rabbit, Licor, 1:800). Cells were then washed 2 times in 1x PBS and then stained with CellTag (CellTag 700, Licor, 1:500) for 1 hour at RT. After a final wash in 1x PBS, the 96-well plate was dried and images were acquired using Li-Cor Odyssey CLx Imaging System.

### I-PpoI mRNA production and transfection

I-PpoI WT and H98A plasmids were generously gifted from Dr. Brian McStay (NUI Galway) and previously characterized^51^. Plasmids were linearized at a NotI site positioned in the polylinker downstream from the I-PpoI ORF and transcribed using MEGAscript T7 kit (Invitrogen) according to the manufacturer’s instructions. I-PpoI mRNA was subsequently polyadenylated using a Poly(A) tailing kit (Invitrogen) according to the manufacturer’s instructions and then precipitated using lithium chloride. SK-Mel28 cells were seeded on poly-l-lysine treated glass coverslips at least 36 hours prior to transfection with the *in vitro* transcribed mRNA using the TransMessenger transfection reagent (Qiagen). One microgram of I-PpoI mRNA and 2μl of Enhancer R were diluted in buffer EC-R to a final volume of 100μl and incubated for 5 minutes at room temperature. Two microliters of TransMessenger transfection reagent was added and further incubated for 10min at room temperature. After addition of 900μl of serum-free medium, the transfection cocktail was added to cells. Following 4 h of incubation, the transfection medium was replaced by full medium, and cells were grown for an additional 2h or 24h prior to further processing.

### RNP transfection

SK-Mel28 cells were grown on coverslips and transiently transfected with ribonucleprotein complexes consisting of purified recombinant Cas9 protein (Truecut Cas9 Protein v2, Invitrogen, #36499) and synthetic guide RNAs (Invitrogen TrueGuide Synthetic sgRNA #35514 or Negative Control, non-targeting 1 #A35526) using Lipofectamine CRISPRMAX Cas9 Transfection Reagent (Invitrogen) according to the manufacturer’s specifications unless otherwise stated. Cells were collected 24 hours post-transfection. The gRNA oligos targeted sequences in the 28S rDNA sequence or a human non-targeting control (Invitrogen, #CMAX00015). Unless otherwise stated, the two rDNA gRNA oligos were pooled in a ratio of 1:1 for each transfection. rDNA guide 1: CGAGAGAACAGCAGGCCCGC; rDNA guide 3: GATTTCCAGGGACGGCGCCT. Cells were collected 24 hours post-transfection. When applicable, Actinomycin D (Act D) was used as a positive control with cells treated at a final concentration of 100ng/ml for 1 hour.

### Laser-induced damage

SK-Mel28 control cells were seeded onto poly-l-lysine treated live-cell imaging glass-bottom 4-well plates at 60,000 cells per well. Cells were then transfected with 800ng Synuclein-GFP or Empty Vector-GFP using Lipofectamine 3000 transfection reagent. Twenty-four hours post-transfection, cells underwent laser-induced damage (LID) using a Zeiss 880 LSM multiphoton microscope outfitted with dual channel BiG (binary GaAsP) detectors and a Coherent Technologies Chameleon titanium-sapphire femtosecond pulsed laser source (for imaging Synuclein-GFP). The Bleaching function is Zen was used to illuminate small, submicron-sized regions within the nucleolus with Chameleon laser tuned to ∼730 nm for 65 or 130µs. There is a ∼4 second time delay required to switch the laser to and from the LID (∼730nm) wavelength. Transmitted light was used to localize the LID pulse to the nucleolus. LID images were analyzed with ImageJ where regions of interest (ROIs) were selected to obtain mean fluorescence values in LID and control ROIs within the nucleolus. The ratio of the signal at each time point from the LID versus the control ROIs was used to calculate the Enrichment Ratio.

### Live-cell nucleolar cap imaging

SK-Mel28 cells were seeded onto poly-l-lysine treated live-cell imaging glass-bottom 4-well plates at 60,000 cells per well. Cells were then transfected with 800ng GFP-Treacle (pcDNA 4TO-Strep-HA-AcGFP-Treacle) using Lipofectamine 3000 transfection reagent. Twenty-four hours post-transfection, cells were transfected with 1µg I-PpoI WT mRNA in serum free media. Four hours after transfected, cells were washed and media was replaced with DMEM Fluorobrite + 10% FBS. Live cell imaging was performed using the Zeiss Celldiscoverer 7 equipped with a 40x oil objective at 37C with 5% CO_2_. Cells were imaged every hour over 15 hours as z-stacks. Images were processed in FIJI and nucleolar caps were hand-counted by investigators blinded to condition.

### IncuCyte proliferation, migration, and invasion assay

SK-Mel28 cells were seeded onto Matrigel treated (80μg/ml) 96-well plates (IncuCyte ImageLock) at 6,000 cells per well. Five hours after seeding, the cells were placed in the IncuCyte for 48 hours for proliferation analysis. At 48 hours, cells were taken out of the IncuCyte and placed in a standard incubator for 72 additional hours to obtain 100% confluence. After 120 hours post-seeding, the confluent cell layer was scratched with an IncuCyte 96-pin wound making tool. An additional 50μl of media (migration) or 50μl of 0.3mg/ml Matrigel (invasion) was added to the wells with scratches. The subsequent movement of cells into the wound was observed and documented with the IncuCyte ZOOM software every 3 hours for 96 hours. The data were exported as the width of the cell-free area. For calculation of the cell migration/invasion distance, the equation the Relative Wound Density was used, where it is a measure (%) of the density of the wound region relative to the density of the cell region:

### 5-EU labeling and immunofluorescence

For 5-EU labeling, the RNP transfection reagents was performed using double the manufacturer’s specifications. 30 minutes prior to fixing cells were incubated with 1 µM 5-Ethynyl Uridine (EU) (Invitrogen #E10345), except for the unlabeled control. Cells were fixed with 4% paraformaldehyde at room temperature (RT) for 10 minutes and subsequently permeabilized with 0.5% Triton X-100 for 10 minutes at RT. Click-iT reaction cocktail (100 Mm Tris-buffered saline, 4 mM CuSO_4_, 100 mM ascorbic acid and 4 µM Alexa Fluor^TM^488 azide (Invitrogen #A10266) was used for detecting EU. Subsequently, samples were incubated at RT with primary antibodies (RNA Polymerase II, rabbit polyclonal, 1:1000, Abcam, ab5131; Treacle, mouse monoclonal, 1:500, Santa Cruz, sc-374536) for 1 hour, 30 minutes with secondary antibody and 5 mg/ml DAPI (Invitrogen #D1206) in a dark humid chamber. Cells were washed three times in PBS between incubation steps, rinsed with water, mounted with Fluoromount-G^TM^ mounting medium (Invitrogen #00-4958-02), and sealed with nail polish. Qualitative image analysis of fluorescence was done using the confocal microscope LSM800 (Zeiss), using the 40x oil immersion objective and ZEN Software (Zeiss). Quantitative analysis was done using CellProfiler^TM^ cell image analysis software. The analysis pipeline first segmented nuclei through the DAPI staining, and subsequently identified nucleoli through an inverted mask of the RNA Polymerase II staining. The nucleolar mask could then be used to measure nucleolar EU intensity per nuclei. For cells treated with Act D or gRNA, cells without nucleolar caps were manually removed for the data analysis. Experiments were repeated three times and at least 100 cells were quantified for each condition per repeat.

### Bulk RNA-sequencing

RNA-sequencing was carried out as previously described^67^. Differential expression results were cross-referenced with antibody-validated nucleolar transcripts previously identified^68^. Nucleolar differentially expressed transcripts were submitted to the Enrichr online gene set enrichment platform^101-103^, and results were visualized with ggplot2. All results were filtered for significance (P < 0.05) and ordered by combined score (log of p value * zscore of deviation from expected rank). Gene sets found to be significant after Benjamini-Hochberg multiple testing correction (P.adj < 0.05) were marked with an asterisk.

### Quantitative RT-PCR

Total RNA was extracted from SK-Mel28 cells (control/KO) using the Qiagen RNeasy Mini Kit (#74104) following the manufacturer’s instructions. RNase-Free DNase Set (Qiagen) was used to remove genomic DNA contaminants. Total RNA was eluted with nuclease-free water. cDNA was synthesized from 500ng RNA with TaqMan Reverse Transcription Reagents Kit (Applied Biosystems). Relative quantification with qRT-PCR was performed by using TaqMan Gene Expression Master Mix (Applied Biosystems) and TaqMan assay reagents (Table 2) on the QuantStudio 3 (Applied Biosystems). Gene expression was normalized to GAPDH expression. SNCA gene expression was used as a positive internal assay control.

### Statistical analysis

Beyond individualized analysis within each assay methodology, all data was processed using GraphPad Prism version 9.0. Data was analyzed using one-way ANOVA, unless stated otherwise, and considered statistically significant if p < 0.05. All data was presented as a mean +/- standard error of the mean (SEM).

## Acknowledgments

All imaging experiments could not have been possible without the help from Stefanie Kaech Petrie and Felice Kelly in the OHSU Advanced Light Microscopy Core. I-PpoI plasmids were graciously gifted by Dr. Brian McStay (NUI Galway). We thank Dr. Pamela Cassidy for the isolation and culturing of the primary melanocytes and PIG1 cells and for the invaluable discussions. Thank you to Dr. Rosie Sears (OHSU) for providing PLA reagents. We thank the Bioimaging Core Facility headed by Dr. Chris Dinant at the Danish Cancer Institute for support. Lastly, a huge thanks to Unni Lab members who helped with the revision process: Valerie Osterberg, Dr. Sydney Boutros, Dr. Carlos Soto Faguas, Elizabeth Rose, Anna Bowman, Elias Wisdom, and Jessica Keating.

Research reported in this publication was supported by the National Institute on Aging of the National Institutes of Health under Award Number F30AG082406. This work was also supported by grants from the NIH (R01NS102227, P30NS061800, P30CA065823). The content is solely the responsibility of the authors and does not necessarily represent the official views of the National Institutes of Health. Research in this publication are also supported by additional external funding sources, including the Danish Cancer Institute, Danish Cancer Society (R302-A17506), Melanoma Research Alliance, Michael J Fox Foundation, and Kuni Foundation.

## Author Contributions

MRA and VU contributed to the conception and design of the study; MRA, KCO, GMC, SNE, CM, PVL, DB, CKM, SNW, DHL, and VU contributed to the acquisition and analysis of data; MRA, SNE, DB, and VU contributed to drafting the text or preparing the figures.

