## Supplemental Figures and Tables for "Alpha-synuclein regulates nucleolar DNA double-strand break repair in melanoma"

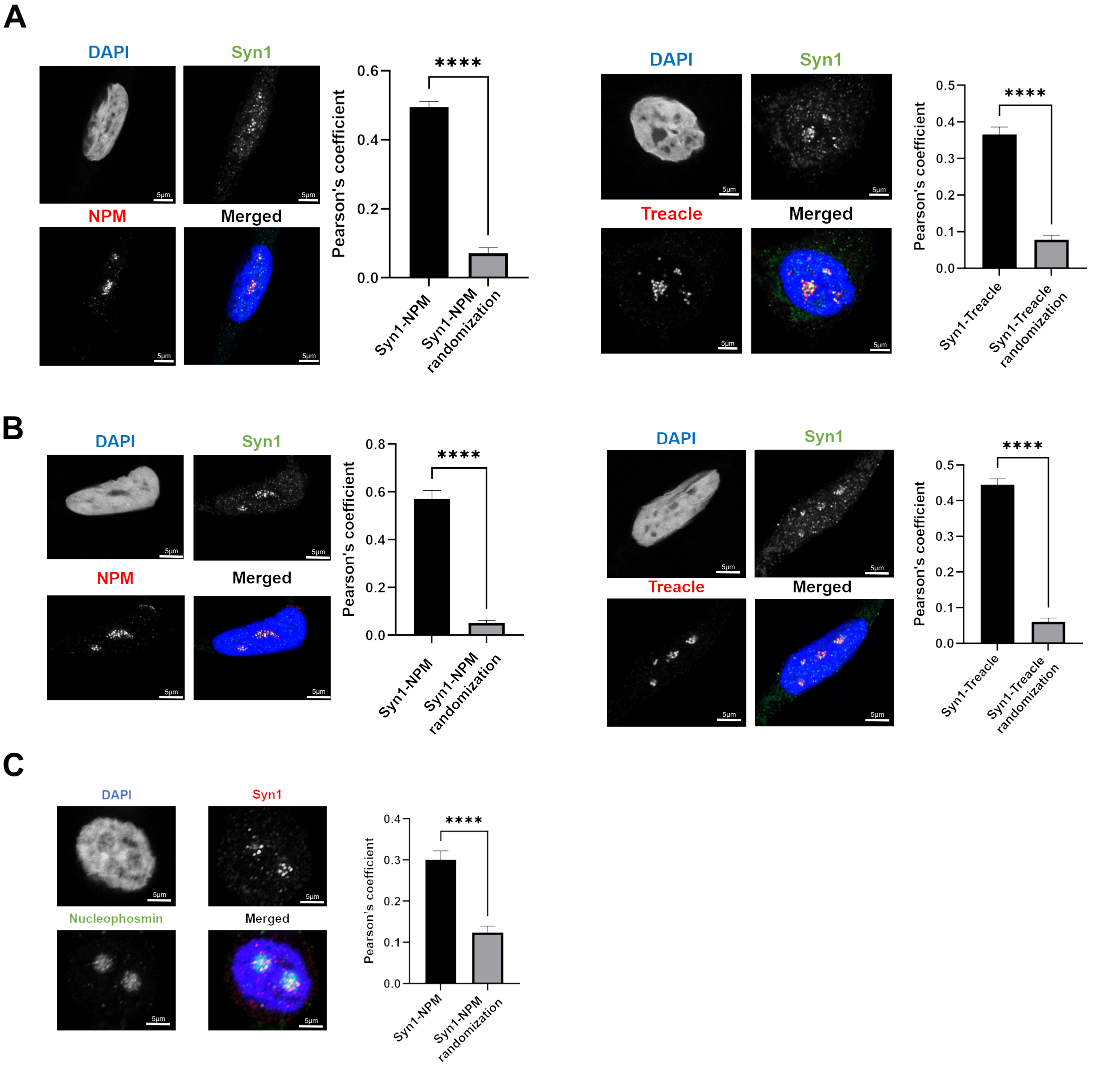


**Supplemental Figure 1. Alpha-synuclein localizes to nucleolar markers in melanoma cells and primary melanocytes.**
A, B, C) A375 (A), PIG1 (B), and primary melanocytes (C) cells were seeded on PDL-coated coverslips and then fixed and stained for alpha-synuclein (Syn1), nucleolar markers (nucleophosmin, treacle), and DAPI. Cells were imaged on the Zeiss 980 confocal microscope with Airyscan and colocalization was analyzed in Imaris software. Error bars represent Standard Error of the Mean (SEM) with quantification from 1 biological replicate. ****p<0.0001 by T-test.

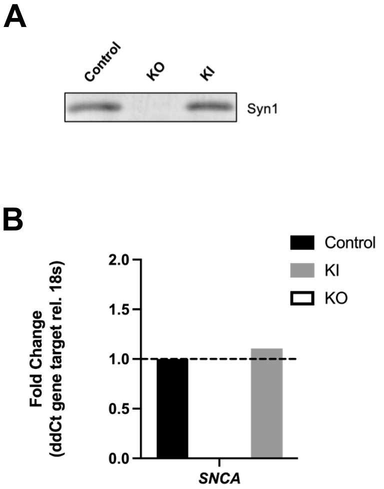


**Supplemental Figure 2. Knock out and reintroduction validation of alpha-synuclein in SK-Mel28 cells.**
A) SK-Mel28 cells (control/KO/KI) were lysed using RIPA buffer and un out on SDS-PAGE and probed for Syn1 and total protein. Representative image of western blot. B) Total RNA was isolated from SK-Mel28 cells (control/KO/KI). Using *SNCA* TaqMan primers, qRT-PCR amplification was determined when normalized to 18S internal control.

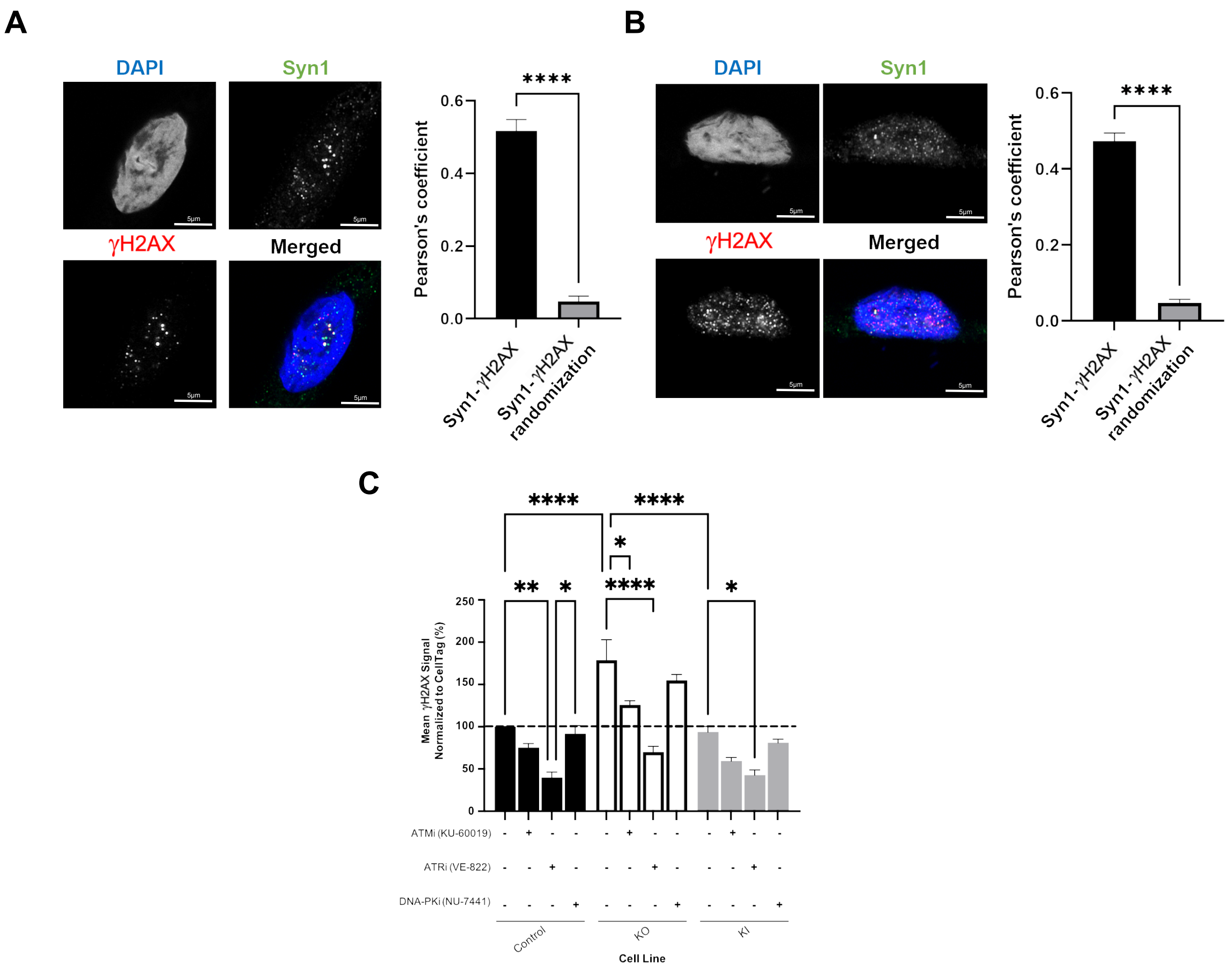


**Supplemental Figure 3. Alpha-synuclein localizes with γH2AX in melanoma cells and primary melanocytes and γH2AX increase is driven by ATM and ATR in alpha-synuclein knockout cells.**
A, B) A375 (A) and PIG1 (B) cells were seeded on PDL-coated coverslips and then fixed and stained for alpha-synuclein (Syn1), DSB marker (γH2AX), and DAPI. Cells were imaged on the Zeiss 980 confocal microscope with Airyscan and colocalization was analyzed in Imaris software. Error bars represent Standard Error of the Mean (SEM) with quantification from 1 biological replicate. ****$p<0.0001$ by T-test. C) SK-Mel28 cells were seeded in a bla ck-welled PDL-coated 96 well plate and treated with DMSO, KU-60019 (10μM), VE-822 (0.1μM), or NU-7441 (1μM) for 24 hours. Cells were processed according to the In-Cell Western manufacturer instructions and stained for γH2AX (800) and CellTag (700). Plates were imaged on the Licor CLx. * $p<0.05$, ** $p<0.01$, **** $p<0.0001$ by ANOVA. Error bars denote SEM. Quantification from 3 biological replicates. Same γH2AX quantification from DMSO condition as Figure 2C as samples were run on the same plate.

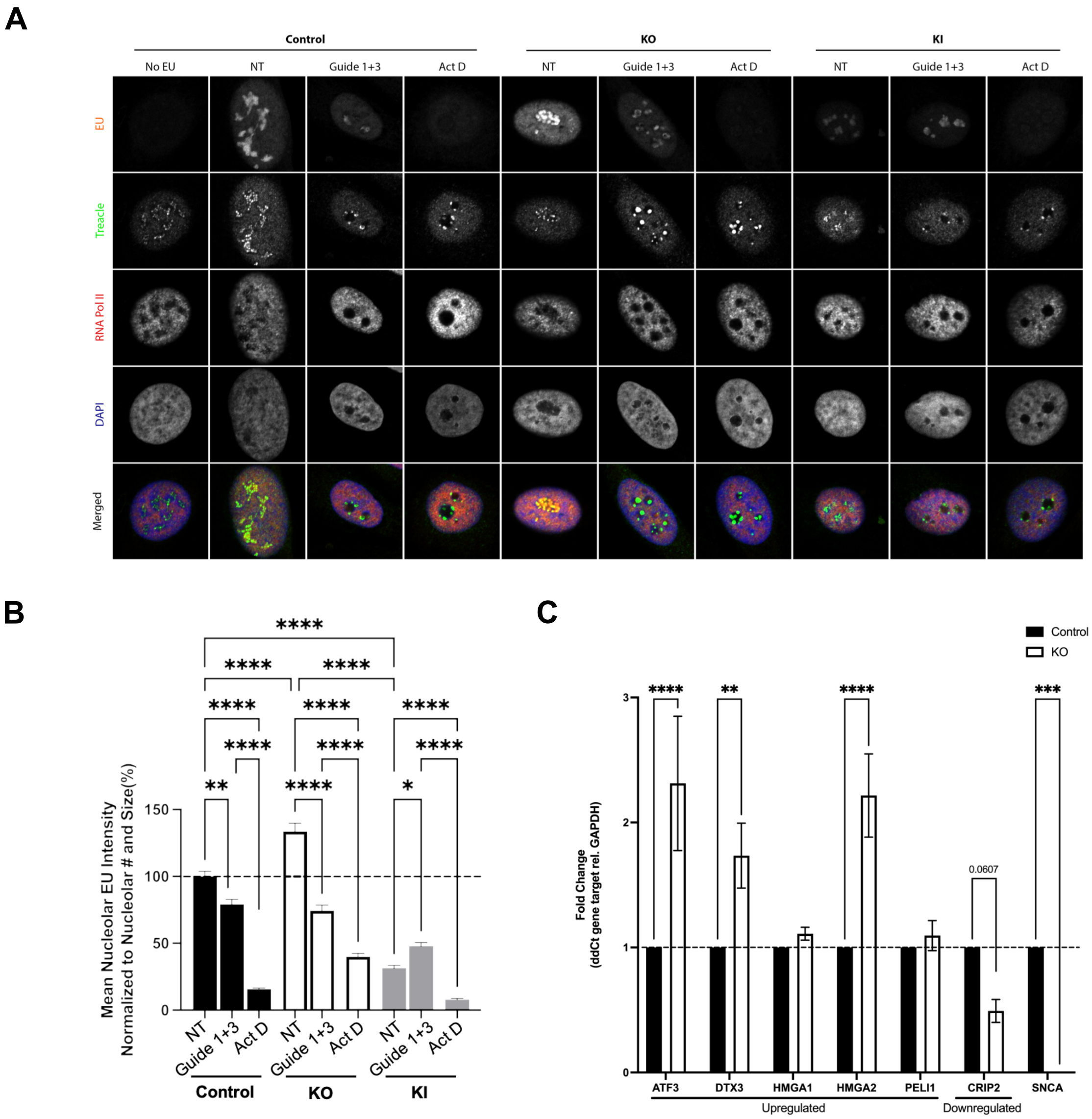


**Supplemental Figure 4. Alpha-synuclein is not involved in nucleolar DDR-mediated transcriptional silencing**
A, B) SK-Mel28 cells (control/KO/KI) were seeded on coverslips and then transfected with Cas9 (TrueCut Cas9 Protein v2, Invitrogen) and guideRNAs that target portions of the 28S unit of rDNA or non-targeting control (NT vs. Guide 1+3). Twenty-four hours after transfection, cells were treated with 5-Ethynyl Uridine before fixing and performing Click-iT reaction to detect EU. Additional staining for treacle, RNA polymerase II, and DAPI were completed. Cells were imaged on the Zeiss LSM800 confocal microscope and analyzed using RNA polymerase II anti-masking. 5EU intensity was measured and normalized to nucleolar pixel number and size. Quantification from 3 biological replicates. * $p<0.05$, ** $p<0.01$, ****$p<0.0001$ by ANOVA. Error bars denote SEM. C) Total RNA from SK-Mel28 cells (control/KO) was isolated and cDNA was prepared using Invitrogen TaqMan Reverse Transcriptase Reagents. Quantitative RT-PCR was performed using TaqMan assay reagents (Table 2) and gene expression levels were normalized to GAPDH. Quantification from 5 biological replicates. ** $p<0.01$, *** $p<0.001$, **** $p<0.0001$ by two-way ANOVA. Error bars denote SEM.

| Gene Name | Nucleolar Location | Fold Change | log2(Fold Change) | PValue | FDR |
|---|---|---|---|---|---|
| ATF3 | Nucleoli | 29.52536727 | 4.8838831 | 1.91E-227 | 2.98E-225 |
| SFRP1 | Nucleoli | 16.07883589 | 4.007091054 | 0 | 0 |
| TBL1X | Nucleoli | 7.685113901 | 2.942066642 | 6.34E-158 | 5.55E-156 |
| DTX3 | Nucleoli | 7.540997575 | 2.914755386 | 2.6E-107 | 1.27E-105 |
| SPOCD1 | Nucleoli | 6.350004811 | 2.666757685 | 4.22E-89 | 1.62E-87 |
| PCDH1 | Nucleoli | 6.316758918 | 2.659184512 | 4.18E-53 | 7.86E-52 |
| DUSP1 | Nucleoli | 6.078912815 | 2.603813327 | 8.99E-166 | 8.96E-164 |
| LURAP1L | Nucleoli | 4.698106043 | 2.232079277 | 1.09E-44 | 1.66E-43 |
| HMGA2 | Nucleoli rim | 3.958533749 | 1.984966151 | 3.57E-78 | 1.19E-76 |
| EN1 | Nucleoli rim | 3.593823411 | 1.845519521 | 1.37E-19 | 8.43E-19 |
| PFKFB4 | Nucleoli | 3.501787008 | 1.808091336 | 2.55E-149 | 2.04E-147 |
| CDCA7L | Nucleoli | 3.485427803 | 1.801335744 | 7.4E-132 | 5.22E-130 |
| BMP7 | Nucleoli | 3.351598842 | 1.744849481 | 9.51E-57 | 1.94E-55 |
| PLOD2 | Nucleoli | 3.283596221 | 1.715276732 | 1.28E-66 | 3.33E-65 |
| PLPP4 | Nucleoli rim | 2.781738245 | 1.475986672 | 9.86E-155 | 8.22E-153 |
| PHLDA1 | Nucleoli rim | 2.685250597 | 1.425056732 | 2.34E-259 | 4.23E-257 |
| PDGFRL | Nucleoli | 2.652722178 | 1.407473589 | 7.97E-71 | 2.26E-69 |
| HIST1H1C | Nucleoli rim | 2.601212023 | 1.379183997 | 9.35E-40 | 1.24E-38 |
| HMGA1 | Nucleoli | 2.58668238 | 1.371102916 | 6.88E-197 | 9.19E-195 |
| SH3TC1 | Nucleoli rim | 2.582615378 | 1.368832803 | 9.98E-26 | 8.1E-25 |
| CSTB | Nucleoli | 2.51382743 | 1.329885615 | 2.49E-187 | 3.06E-185 |
| KLF6 | Fibrillar center | 2.460415276 | 1.298901838 | 5.08E-161 | 4.75E-159 |
| DLG3 | Nucleoli | 2.448057714 | 1.291637571 | 5.95E-24 | 4.5E-23 |
| FAM198B | Fibrillar center | 2.445531037 | 1.290147775 | 3.04E-25 | 2.41E-24 |
| KCNC4 | Nucleoli | 2.439896714 | 1.286820077 | 1.73E-44 | 2.62E-43 |
| OSMR | Nucleoli | 2.300357658 | 1.201858188 | 2.26E-104 | 1.05E-102 |
| PELI1 | Nucleoli | 2.272113094 | 1.184034646 | 4.11E-49 | 7.03E-48 |
| CTSB | Nucleoli | 2.23062657 | 1.157449012 | 1.78E-169 | 1.84E-167 |
| FADS3 | Fibrillar center | 2.20816123 | 1.142845515 | 7.9E-52 | 1.46E-50 |
| DDX41 | Nucleoli | 2.156596089 | 1.108755998 | 6E-85 | 2.19E-83 |
| B3GNT5 | Nucleoli | 2.131456723 | 1.091839763 | 9.73E-18 | 5.46E-17 |
| SMPDL3A | Nucleoli | 2.108575404 | 1.076268615 | 1.06E-32 | 1.12E-31 |
| SLC6A15 | Nucleoli | 2.100790212 | 1.070932099 | 1.73E-47 | 2.85E-46 |
| CCDC59 | Nucleoli rim | 2.091777836 | 1.064729633 | 1.24E-36 | 1.49E-35 |
| RNFT1 | Nucleoli | 2.03805413 | 1.027192369 | 1.14E-19 | 7.04E-19 |
| PELO | Fibrillar center | 2.016925559 | 1.012157837 | 2.25E-64 | 5.49E-63 |
| ATP6AP1L | Nucleoli | 2.013390254 | 1.009626836 | 2.55E-49 | 4.4E-48 |
| ICK | Fibrillar center | -2.000231199 | -1.000166765 | 1.08E-21 | 7.34E-21 |
| DPH6 | Nucleoli rim | -2.005977839 | -1.004305668 | 3.59E-16 | 1.85E-15 |
| ZNF397 | Nucleoli | -2.013859384 | -1.009962952 | 5.51E-16 | 2.82E-15 |
| OSCP1 | Nucleoli | -2.024484901 | -1.017554883 | 3.04E-13 | 1.32E-12 |
| FOXJ2 | Fibrillar center | -2.027796062 | -1.019912566 | 3.27E-47 | 5.35E-46 |
| ZNF689 | Fibrillar center | -2.040976132 | -1.029259311 | 3.71E-160 | 3.4E-158 |
| UBR3 | Nucleoli | -2.079282131 | -1.056085526 | 8.36E-73 | 2.47E-71 |
| LZTS1 | Nucleoli | -2.100734509 | -1.070893846 | 3.34E-134 | 2.42E-132 |
| MYL5 | Nucleoli | -2.112441599 | -1.078911457 | 1.02E-13 | 4.56E-13 |
| AKAP11 | Nucleoli | -2.154422695 | -1.107301332 | 4.51E-57 | 9.27E-56 |
| BRWD1 | Nucleoli | -2.158029608 | -1.109714658 | 2.45E-66 | 6.31E-65 |
| ZNF33B | Fibrillar center | -2.198838417 | -1.136741591 | 8.08E-33 | 8.57E-32 |
| ZBTB43 | Nucleoli | -2.239812724 | -1.16337811 | 1.03E-88 | 3.96E-87 |
| TTC28 | Nucleoli | -2.242272354 | -1.164961523 | 1.65E-77 | 5.44E-76 |
| TAF4B | Fibrillar center | -2.378460438 | -1.250028029 | 1.66E-49 | 2.91E-48 |
| TMOD2 | Fibrillar center | -2.463846408 | -1.300912323 | 1.59E-60 | 3.5E-59 |
| RBM43 | Nucleoli | -2.499687886 | -1.321747969 | 1.03E-29 | 9.78E-29 |
| ABCC4 | Nucleoli | -2.539678916 | -1.344646113 | 1.9E-105 | 8.93E-104 |
| IQSEC1 | Nucleoli rim | -2.559626688 | -1.355933414 | 3.08E-71 | 8.81E-70 |
| SPIN4 | Nucleoli | -2.76010701 | -1.464724201 | 1.39E-21 | 9.41E-21 |
| MYL9 | Fibrillar center | -2.840665986 | -1.506229205 | 9.12E-53 | 1.7E-51 |
| POP1 | Nucleoli | -2.840967431 | -1.506382293 | 3.57E-109 | 1.79E-107 |
| JRK | Nucleoli | -2.960246116 | -1.565717127 | 1.26E-79 | 4.24E-78 |
| ZNF419 | Fibrillar center | -3.137808254 | -1.649757194 | 7.28E-23 | 5.24E-22 |
| STOX1 | Fibrillar center | -3.498043645 | -1.80654829 | 5.71E-29 | 5.27E-28 |
| BCAM | Fibrillar center | -5.401784718 | -2.433436144 | 3.55E-59 | 7.67E-58 |
| CRIP2 | Nucleoli | -8.074263539 | -3.013330676 | 1.58E-154 | 1.31E-152 |

**Table 1. Differentially expressed nucleolar transcripts in alpha-synuclein knockout cells.** Total RNA was extracted from SK-Mel28 control and KO cells and sent for RNA-sequencing analysis at Indiana University. Differentially expressed gene transcripts in KO cells compared to control were identified. These were cross-referenced to over 500 nucleolar-specific genes. 64 genes were identified.

| Gene of Interest | TaqMan Assay |
|---|---|
| ATF3 | Hs00231069_m1 |
| DTX3 | Hs01595350_m1 |
| HMGA1 | Hs00852949_g1 |
| HMGA2 | Hs04397751_m1 |
| PELI1 | Hs00900505_m1 |
| CRIP2 | Hs00373842_g1 |
| SNCA | Hs00240906_m1 |
| GAPDH | Hs02786624_g1 |
| 18S | Hs99999901_s1 |

**Table 2. TaqMan assay probes**
